# MAGI1 inhibits the AMOTL2/p38 stress pathway and prevents luminal breast tumorigenesis

**DOI:** 10.1101/2020.06.13.149724

**Authors:** Diala Kantar, Emilie Bousquet Mur, Maicol Mancini, Vera Slaninova, Yezza Ben Salah, Luca Costa, Elodie Forest, Patrice Lassus, Charles Géminard, Florence Boissière-Michot, Béatrice Orsetti, Charles Theillet, Jacques Colinge, Christine Benistant, Antonio Maraver, Lisa Heron-Milhavet, Alexandre Djiane

## Abstract

Alterations to cell polarization or to intercellular junctions are often associated with epithelial cancer progression, including breast cancers (BCa). We show here that the loss of the junctional scaffold protein MAGI1 is associated with bad prognosis in luminal BCa, and promotes tumorigenesis. E-cadherin and the actin binding scaffold AMOTL2 accumulate in *MAGI1* deficient cells which are subjected to increased stiffness. These alterations are associated with low YAP activity, the terminal Hippo-pathway effector, but with an elevated ROCK and p38 Stress Activated Protein Kinase activities. Blocking ROCK prevented p38 activation, suggesting that MAGI1 limits p38 activity in part through releasing actin strength. Importantly, the increased tumorigenicity of *MAGI1* deficient cells is rescued in the absence of AMOTL2 or after inhibition of p38, demonstrating that MAGI1 acts as a tumor-suppressor in luminal BCa by inhibiting an AMOTL2/p38 stress pathway.

## INTRODUCTION

Breast cancers (BCa) arise from mammary epithelial cells, and show tremendous diversity. Based on transcriptome, BCa are typically divided into five main subtypes: luminal A, luminal B, HER2-enriched, normal-like, and basal (reviewed in ^1 2^, partly overlapping with previous classifications (such as ER/PR/HER2 receptors expression status). Basal tumors, overlapping strongly with triple-negative breast cancers, are the most aggressive and present the worst prognosis. However, even though luminal A and B subtypes (typically ER/PR positives) usually respond well to current treatments, when “luminal” tumors relapse, they become very difficult to treat with very poor outcome.

The apico-basal (A/B) polarity of mammary epithelial cells is established and maintained by the asymmetric segregation of evolutionarily conserved protein complexes ^3^. A/B polarity governs the correct position of different intercellular junctions that ensure the integrity of the epithelial sheet. Adherens Junctions (AJs) constitute apical adhesive structures mediated by the homophilic trans-interactions of E-cadherin/catenin complexes ^4 5^. At the very apical part of epithelial cells, Tight junctions (TJs) ensure the impermeability of tissues in part through the action of Claudins/Zonula Occludens proteins complexes ^6^. The transmembrane proteins of these different junctions (E-cadherin, Claudins...) are coupled and anchored to the actin cytoskeleton via adaptor scaffold proteins, thus coupling adhesion and cell contacts to actin dynamics. As such AJs represent mechano-sensory structures, where tensile or compressive forces, arising from internal or external sources, will be sensed to promote short term responses such as changes in adhesion and actomyosin dynamics, but also long term responses to control cell density, in particular through the mechano-sensitive Hippo pathway^7,8^. In epithelial cells, the compressive forces exerted by the cortical actin cytoskeleton or by cell packing, sensed at AJs, antagonize the proliferative and anti-apoptotic effects of YAP/TAZ in part through α-catenin conformational change, or through the release of Merlin/NF2 ^9–14^. Other signaling pathways such as β-catenin, ERK, or JNK/SAPK stress pathways can also be influenced by polarity and forces sensed at junctions^11,15–18^. BCa are derived from transformed mammary epithelia where frequent alterations in TJs and/or AJs composition and structure, and the associated abnormal mechano-sensory responses have been linked to cell dysfunction such as uncontrolled cell proliferation and metastasis ^19,20^.

At apical junctions, cortical scaffold proteins such as MAGI (MAGUKs with Inverted domain structure), assemble large and dynamic molecular complexes. In humans, among the three MAGI proteins (MAGI1-3), MAGI1 is the most widely expressed and contains several protein-protein interaction domains including six PDZ and two WW domains ^21^ mediating interactions with TJ proteins ^22^. In *Drosophila*, we have demonstrated that Magi, the sole fly MAGI homolog, is required for E-cadherin belt integrity and AJ dynamics ultimately restricting cell numbers in the developing eye ^23,24^. The regulation of Cadherin complexes by MAGIs, is evolutionarily conserved from C. elegans ^25^ to vertebrates, and in particular vertebrate endothelial cells where MAGI1 links VE-cadherin complexes with the underlying cortical actin cytoskeleton ^26^. A tumor suppressive role of MAGI1 (reviewed by Feng et al.^27^) is suggested by the correlation between low *MAGI1* expression and poor prognosis in various cancers, such as hepatocellular carcinoma ^28^. The anti-tumoral activity of MAGI1 is further supported by its ability to bind the tumor-suppressor PTEN ^29–31^, the fact that it is often targeted by viral oncoproteins ^32,33^, and by the observation that in colorectal cancer cells, MAGI1 was upregulated in response to cyclooxygenase-2 inhibitor and prevented metastasis ^34^. Thus it appeared that the apical junction-localized MAGI1 scaffold protein participates in multiple complexes to fine-tune adhesion and signaling, and may therefore be considered as a tumor suppressor.

The angiomotin (AMOT) family of membrane-associated scaffold proteins is composed of three members: AMOT, AMOTL1, and AMOTL2, playing important roles in the regulation of intercellular junctions ^35^. In endothelial cells and in zebrafish developing embryos, AMOTL2 has been shown to link cadherins and the actin cytoskeleton. In particular, AMOTL2 is required for the maintenance of tension at the level of cadherin junctions to properly shape strong blood vessels ^26,36^. Moreover, AMOTs are also known to control the Hippo tumor-suppressor signaling pathway. Canonical Hippo pathway, through sequential activation of the Hippo/MST and Wts/LATS kinases, promotes the phosphorylation and cytoplasmic retention of the transcriptional co-activators Yki/YAP/TAZ (reviewed in ^37^). Specific PPxY motifs found in AMOTs interact with the WW domains of YAP and TAZ ^38^ and AMOTs act thus as negative regulators of YAP/TAZ mediated transcription by trapping YAP ^38,39^ and TAZ ^40^ in the cytoplasm. Recent studies identified interactions between MAGI1 and AMOTs, in particular in the regulation of intercellular junctions ^41 42,43^.

We report here that low levels of *MAGI1* are associated with bad prognosis in luminal ER+ BCa. Impairing MAGI1 expression in luminal BCa cells promoted their growth in 2D, and their ability to grow in low attachment conditions in-vitro and in subcutaneously grafted nude mice. We further document that *MAGI1* deficient cells accumulate specific junctional proteins, including E-cadherin and AMOTL2. Mechanistically, we show that all *MAGI1* deficient phenotypes are suppressed by down-regulating *AMOTL2* suggesting that AMOTL2 and its effects on junctions is the primary cause of the increased tumorigenicity after MAGI1 loss. Consistent with these observations, we could further show that *MAGI1* deficient luminal BCa cells experience higher stiffness as evidenced by increased Young’s modulus, and ROCK activity as evidenced by increased ROCK-specific Ser19 phosphorylation of the regulatory Myosin Light Chain 2. They activate the p38 stress signaling pathway while YAP activity is antagonized. Finally, we provide evidence that releasing actin cytoskeletal strength, or impairing p38 activity can revert the effects of MAGI1 loss, supporting a model by which, in response to MAGI1 loss, elevated AMOTL2/E-cadherin and junction dysfunction, together with actin cytoskeletal tension activate the p38 stress pathway fueling tumorigenicity.

## RESULTS

### The loss of MAGI1 enhances tumorigenicity of luminal breast cancer (BCa) cells

In order to study the role of the apical scaffold MAGI1 during breast carcinogenesis, we first analyzed the expression of MAGI1 by western blot in both luminal and basal BCa cell lines: MAGI1 expression was restricted to luminal ER+ lineages (T47D and MCF7 luminal A, and to a lesser extent in ZR75 luminal B), while no expression could be detected in basal ER-lineages (immortalized MCF10A and triple negative MDA-MB-468 and BT549; Supplementary Fig. S1A). Moreover, public database mining (http://www.kmplot) ^44^ and recent studies ^45^, indicated that low MAGI1 expression levels were associated with worse prognosis in relapse-free survival for BCa patients, but only in ER+ (mainly luminal) molecular BCa subtypes (Kaplan Meier curve; Supplementary Fig. S1B).

We thus decided to investigate the functional role of MAGI1 and the consequence of MAGI1 knockdown in luminal BCa, and generated MCF7 and T47D cell lines in which *MAGI1* was targeted by constitutive shRNA. Two independent shRNA constructs targeting different parts of the *MAGI1* transcript were used, *shMAGI1(3-1)* and *shMAGI1(1-1)*, which led respectively to more than 90% and around 60% MAGI1 knockdown at the protein level as shown by western blot and immunofluorescence analyses (Supplementary Fig. S1D&E). The knockdown was specific for MAGI1 and did not induce compensatory up-regulation of MAGI2 and MAGI3, the two remaining MAGI family members (Supplementary Fig. S1D&F). *shMAGI1(3-1)* was the most potent resulting in almost knock-out like down-regulation, and was therefore used in all subsequent studies and referred to as *shMAGI1* thereafter. It should be noted that in the subsequent analyses of MAGI1 function similar effects were obtained using the independent *shMAGI1(1-1)* ruling out non-specific phenotypes of the *shMAGI(3-1)* (Supplementary Fig. S2A-F).

First, we studied the effect of *MAGI1* knockdown on cell numbers and proliferation. *MAGI1* depletion increased 2D cell growth of MCF7 cells (20% to 25% increase at day 4 and day 7) as assayed by MTT cell growth assay (Figure 1A). At day 7, it was associated with an increase in the proportion of cells in the S phase of the cell cycle from 41% in control MCF7*shLuc* to 54% in MCF7*shMAGI1* cells as assessed by BrdU incorporation (Figure 1B) with no discernable change in apoptotic cells (see Figure legend 1B). More importantly, in anchorage-independent soft agar cell growth conditions, MCF7*shMAGI1* cell lines formed circa twice as many colonies compared to *shLuc* controls (Figure 1C). Similarly, MCF7*shMAGI1* cells showed enhanced clonal mammosphere capacities which were correlated with an expanded area occupied by the spheres (60% increase; Figure 1D). This increase in mammosphere formation is associated with a slight increase in the proportion of cells presenting stemness markers (to circa 4% of total cells) as evidenced by increase in CD44High/CD24Low (1.7x) and ALDH activity (2.6x). Then, we studied the effect of MAGI1 knock-down on cell migration and could not detect any effect on collective cell migration (scratch wound assay), and on invasion (boyden chamber assay; Supplementary Fig. S3A-C). Importantly, similar results were obtained in T47D, a second luminal A BCa cell line, in which MAGI1 knockdown (*shMAGI1*) led to elevated 2D and anchorage-independent cell growth (Supplementary Fig. S4A,B&G). Taken together, these results show that the loss of MAGI1 in luminal BCa cells promotes cancer cell proliferation and anchorage-independent growth, but not migration/invasion.

**Figure 1.**
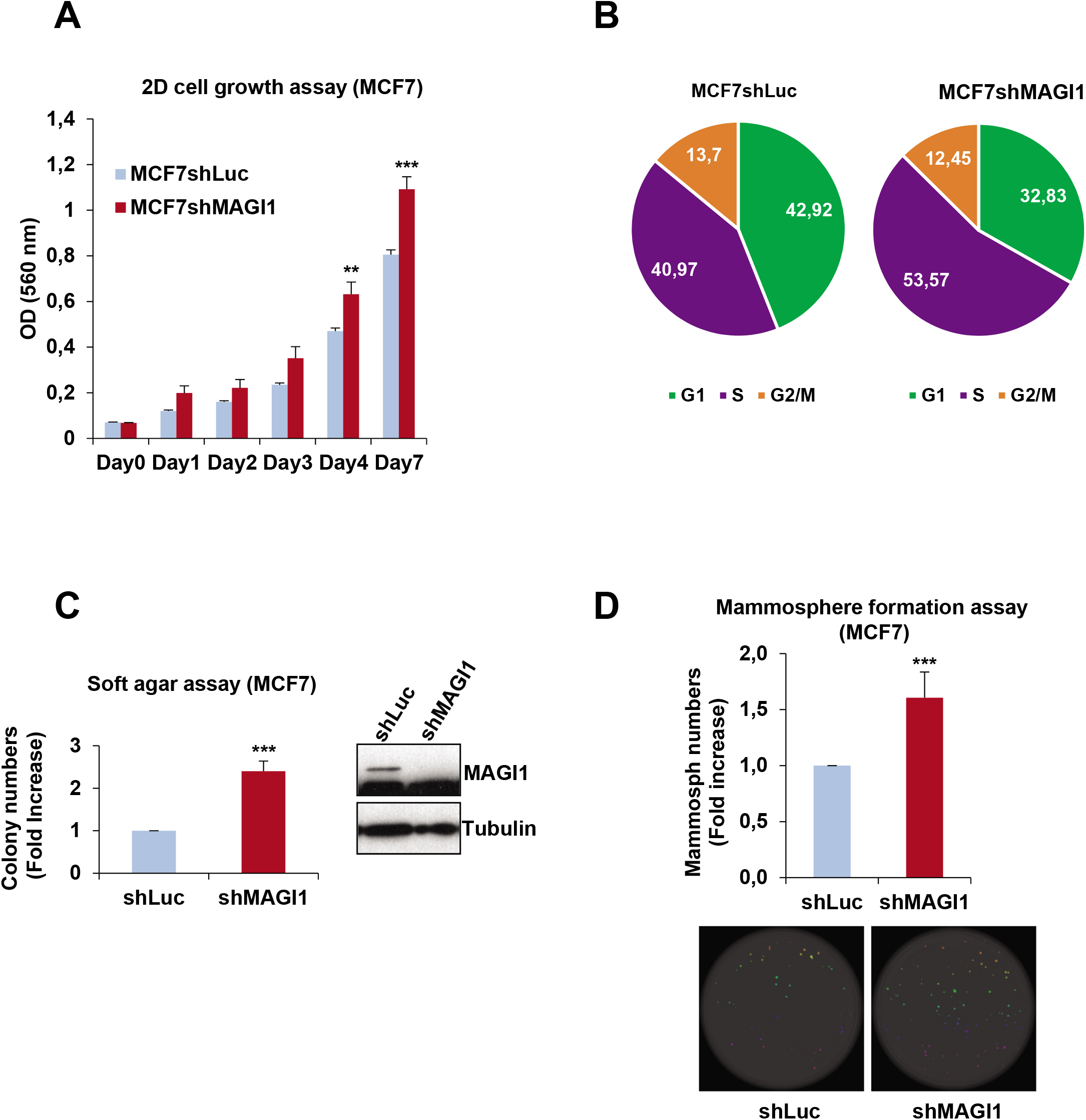
MAGI1 impairment induces tumorigenic phenotypes in epithelial cells. (A) MTT assay (OD 560 nm) representing 2D cell growth of MCF7*shMAGI1* as compared to MCF7*shLuc* cells. Bars represent mean ± Standard Deviation (SD; n=10 wells as replicates) of a representative experiment (out of 3). Unpaired two-tailed Student’s t-test; ** p < 0.01; *** p < 0.001. (B) Cell cycle phases of MCF7*shLuc* and MCF7*shMAGI1* cells assessed after BrdU incorporation and analyzed by flow cytometry, showing an increased proportion of cells in S phase at the expense of G0/G1 after *MAGI1* invalidation. The pie chart also showed no major difference in the percentage of apoptotic cells in SubG1 phase (around 2 % in *shLuc* and 1 % in *shMAGI1* cells). (C) Left: quantification of colony numbers of MCF7*shMAGI1* cells grown in anchorage independent conditions (soft agar assay) and represented as fold increase compared to MCF7*shLuc* cells. Data are presented as the means ± SD (n=3). Unpaired two-tailed Student’s t-test; *** p < 0.001. Right: western blot analysis of whole protein extracts issued from the cells used in the soft agar assays and showing the relative amounts of MAGI1 in the different cell lines; Tubulin was used as a loading control. Uncropped blots can be found in the Supplementary Information. (D) Top: quantification of mammospheres represented as total area issued from MCF7*shMAGI1* cells and shown as fold increase compared to MCF7*shLuc* cells, starting from 200 cells. Data are presented as the means ± SD (n=4). Unpaired two-tailed Student’s t-test; *** p < 0.001. Bottom: images of mammosphere formation assay showing a representative experiment.

Significantly, when MCF7*shMAGI1* cells were injected subcutaneously in nude mice, they grew as a tumor mass much more rapidly and extensively than control MCF7*shLuc* cells (Figure 2A). This dramatic increase in tumor growth was not due to obvious changes in the tumor microenvironment and comparable density of immune cells, fibroblasts, and vascular cells infiltrates were observed in shMAGI1 tumors compared to controls as reflected by CD45, a-SMA, and CD31 staining respectively (Supplementary Fig. S5A-J). Importantly, higher Ki67 reflecting increased number of mitotic cells was observed in the MCF7shMAGI1 tumors (Figure 2B-D) and among the proliferative tumor cells, the Ki67 staining was more intense (Figure 2E), showing that indeed MAGI1-deficient cells proliferate more, and suggesting that the increased tumor growth upon MAGI1 depletion is primarily due to increased tumor cells burden. Even though BCa was the prime focus of this study, the effects of *MAGI1* knockdown appeared not restricted to mammary cells, and similar observations on anchorage-free growth in soft agar, and orthotopic tumor growth in nude mice were also observed using HCT116 colon cancer cells (Supplementary Fig. S4E-G) extending earlier reports ^34^. Altogether, our results show that the loss of MAGI1 in luminal A BCa cells enhances their tumorigenic behaviors.

**Figure 2.**
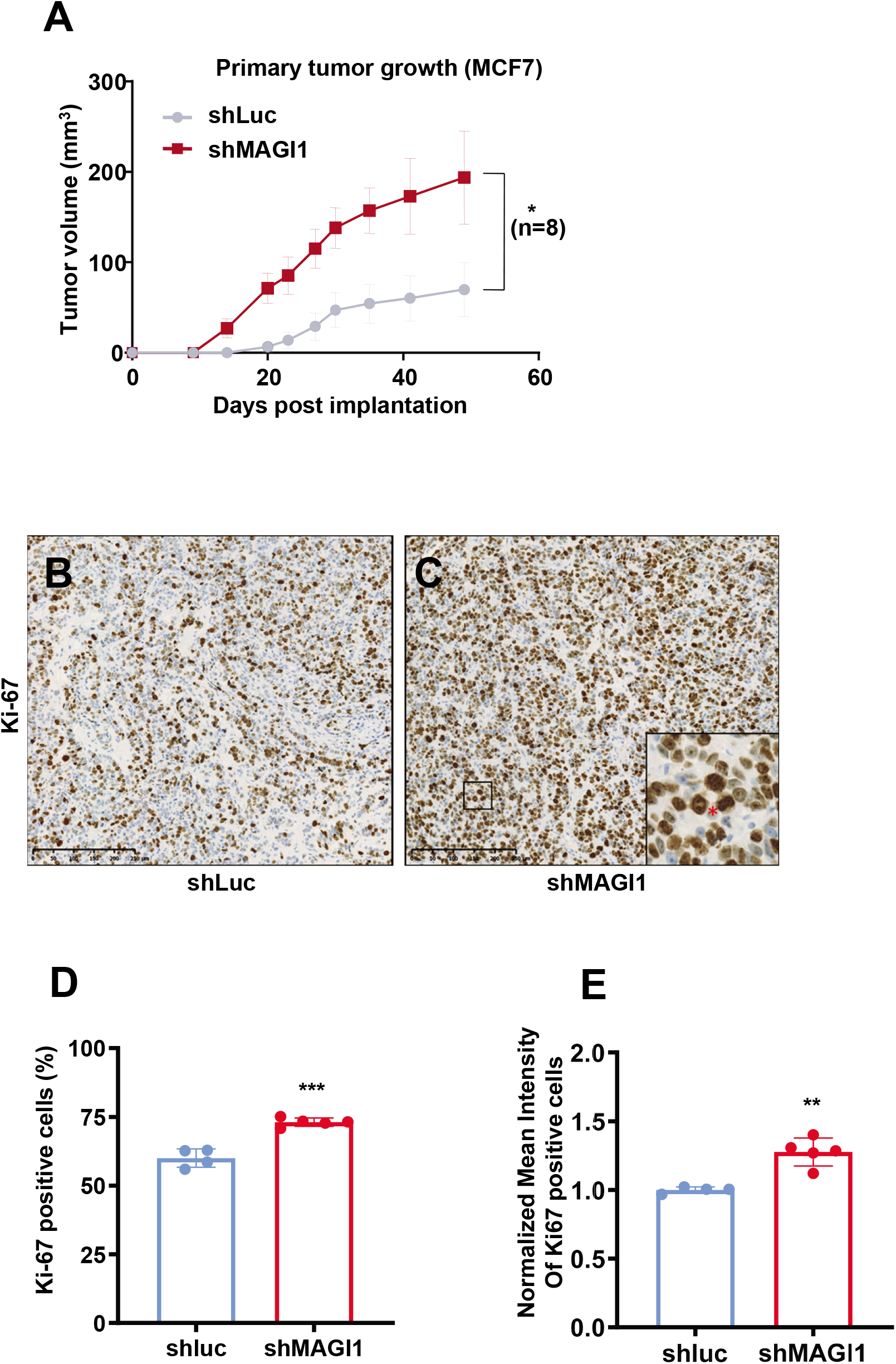
The loss of MAGI1 in primary tumors induces tumor cell proliferation. (A) Primary tumor growth of MCF7*shLuc* and MCF7*shMAGI1* cells injected subcutaneously in nude mice. Primary tumor growth was assessed by measuring the tumor volume over time until the tumors were too big and the mice had to be euthanized. Bars correspond to the mean ± Standard Error to the Mean (SEM; n=8 mice per group). Unpaired two-tailed Student’s t-test; * p < 0.05. (B-C) Representative immunohistochemical staining of Ki67 in primary tumors obtained after subcutaneous engraftment of either MCF7*shLuc* (B) or MCF7*shMAG1* cells (C). Brown staining indicates positive immunoreactivity. Scale bar=250 μM. In the magnified area, the red * highlights an increased number of mitotic cells in the MCF7*shMAGI1*-derived primary tumor. (D-E) Quantification of the immunohistochemical analysis in B&C, showing the percentage of primary tumor cells stained positively for Ki67 (D) and the normalized mean intensity of Ki67 positive cells (E) in MCF7*shMAGI1* and MCF7*shLuc* primary tumors. Bars correspond to the mean ± SD (n=4 for *shLuc* and n=5 for *shMAGI1*). Unpaired two-tailed Student’s t-test; ** p < 0.01 and *** p < 0.001.

### The loss of MAGI1 promotes the accumulation of epithelial junctions components

In order to further investigate the role of MAGI1 in breast tissue, we sought to determine its localization in mammary epithelial cells. Immunohistochemistry on BCa patient tissue micro-array revealed that in normal breast cells, MAGI1 was expressed only in the luminal epithelial cells and not in the underlying basal myo-epithelial layer, consistent with MAGI1 expression exclusively in luminal-type BCa cell lines (Supplementary Fig. S1A). At the sub-cellular level, MAGI1 was localized at the apical pole of luminal breast cells (Supplementary Fig. S1C, arrows). Using MCF7 cancer cells, immortalized hMEC (human mammary epithelial cells), and polarized canine MDCK cells, we further confirmed by immunofluorescence that MAGI1 localized near the plasma membrane, overlapping with junction components such as E-cadherin (AJ), ZO1, and Claudin3 (TJ), suggesting that MAGI1 is a TJ and/or AJ resident protein (Figure 3A, arrows and data not shown).

**Figure 3.**
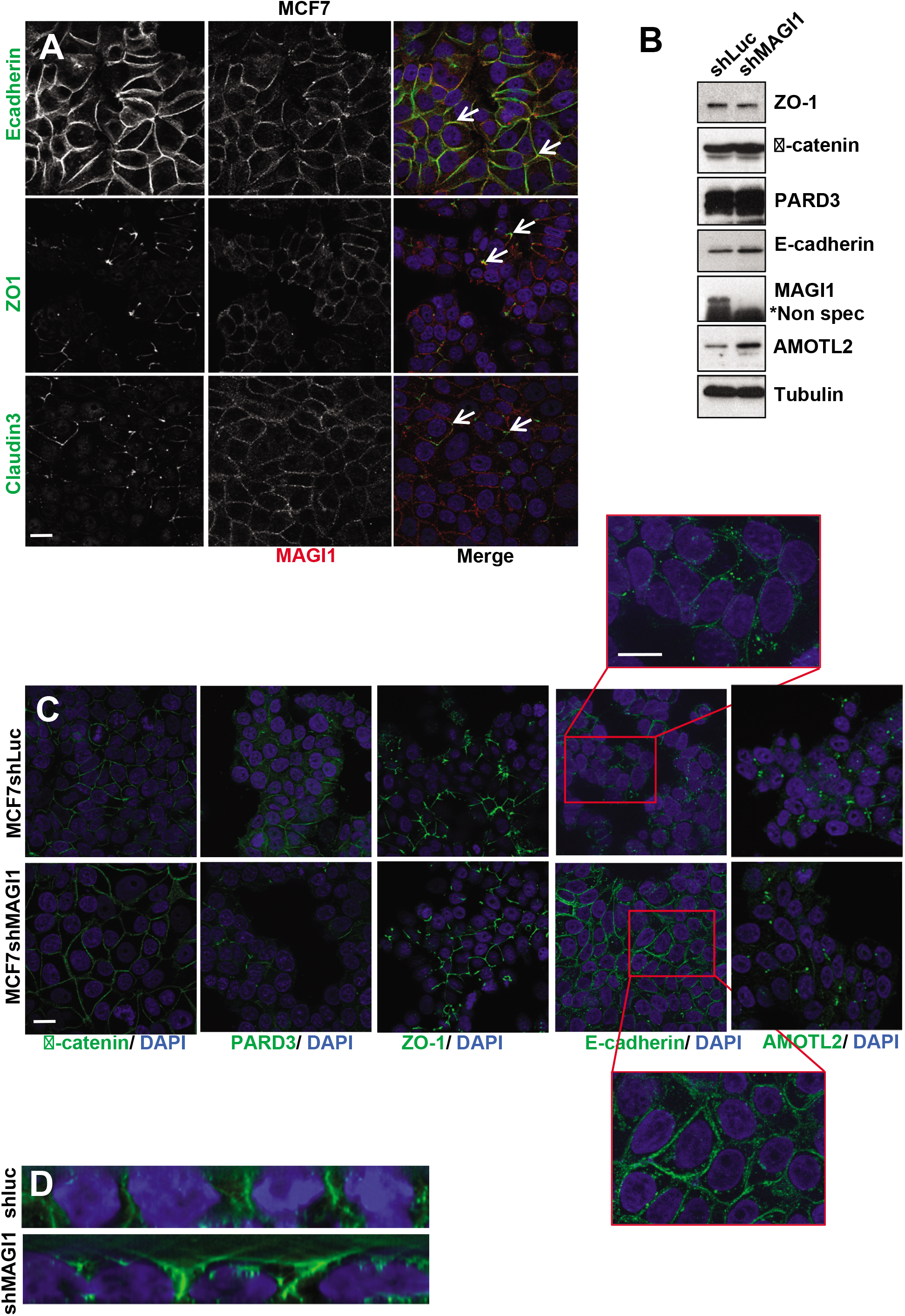
The loss of MAGI1 affects E-cadherin levels and localization in MCF7 cells. (A) Representative immunofluorescence experiments performed in MCF7 cells monitoring MAGI1 localization (red) compared to E-cadherin, ZO1, and Claudin3 (green). DAPI (blue) was used to stain DNA and the nuclei. White arrows indicate staining overlap. Scale bar=10 μm. (B) Western blot analysis of whole protein extracts (n=4) monitoring the expression the junctional components Claudin-1, ZO-1, β-catenin, PARD3, E-cadherin and AMOTL2 in MCF7*shMAGI1* cell lines compared to MCF7*shLuc*. Tubulin was used as a loading control. Note the increase in E-cadherin levels. Western blot experiments have been repeated at least four times. Protein expression levels were quantified as compared to Tubulin. Uncropped blots can be found in the Supplementary Information. (C) Representative immunofluorescence experiments (n=3) were performed on MCF7*shMAGI1* compared to MCF7*shLuc* cells monitoring the localization of β-catenin, PARD3, Claudin-3, ZO-1 and E-cadherin and AMOTL2 (green). DAPI (blue) was used to stain DNA and the nuclei. Scale bar=10 μm. Note the slight sub-cortical accumulation of E-cadherin shown in the high magnification images. (D) Representative E-cadherin staining (green) of MCF7*shLuc* and MCF7*shMAGI1* shown through the Z axis. DAPI (blue) was used to stain DNA and the nuclei.

The localization of MAGI1 at apical junctions prompted us to explore whether MAGI1 could control their biology. Performing western-blot analyses on whole protein extracts, the loss of MAGI1 did not affect the total protein abundance of the typical TJs component ZO-1 nor of the AJs component PARD3 (Figure 3B). However, in MCF7*shMAGI1* we observed increases in the junctional proteins β-catenin (x1.25), E-cadherin (x1.6) and AMOTL2 (x1.8) (Figure 3B). Consistently, performing immunofluorescence on fixed MCF7*shMAGI1* cells, the membrane levels and localization of ZO-1 and PARD3 were unchanged, while E-cadherin staining was increased, and less tightly restricted to the plasma membrane, and extending more basally than controls (Figure 3C&D). The altered distribution of E-cadherin in *shMAGI1* cells is reminiscent of the role of Magi in *Drosophila* where we showed that it controls E-cadherin belt at AJs in epithelial cells of the developing eye ^24^.

In epithelial cells, the increase in E-cadherin material is often associated with increased junctional strength and, through the linkage to the underlying actin, to increases in tension and overall compressive forces (reviewed by ^7^). While we did not observe obvious changes in the intensity or morphology of F-actin stained with phalloidin upon MAGI1 knockdown (Supplementary Fig. S6A), MCF7*shMAGI1* cells exhibited behaviors compatible with increased compression. First, when cultured as spheroids, MCF7*shMAGI1* cells grew as round masses of cells, rounder than MCF7*shLuc* controls with circa 30% smaller perimeter (Figure 4A&B) and better circularity (0.95 vs 0.62 respectively; Figure 4C), a sign that MCF7*shMAGI1* cells aggregates were more compact, even though composed of the same number of cells as MCF7*shLuc* controls. It is noteworthy that MCF7*shMAGI1* spheroids grew more (more cells) than controls reminiscent of the effect of MAGI1 knock-down in 2D and anchorage-independent cell growth assays (Figure 4D). This compaction was also observed in T47D *shMAGI1* cells (Supplementary Fig. S4C&D). Second, using Atomic Force Microscopy (AFM), we showed that MCF7*shMAGI1* cells were stiffer than controls (higher Young’s modulus factor; Figure 4E), strongly suggesting that MCF7*shMAGI1* cells may have a stronger cytoskeleton and might experience higher internal pressure. Since AFM was performed on small MCF7 cells clusters, our data indicate that the increased compressive forces were generated from within the cells, likely through increased actin contractility. Indeed, elevated ROCK activity was detected in MAGI1 deficient cells as shown by the increase in ROCK-specific serine 19 phosphorylation of the regulatory myosin light chain MLC2 (Figure 4F&G), suggesting that the actin cytoskeleton was under higher tension. Consistently, more actin-associated phosphorylated MLC2 (Ser19) positive foci were detected in MCF7*shMAGI1* cells compared to MCF7*shLuc* cells (Supplementary Fig. S6A&B). Furthermore, treating MCF7*shMAGI1* cells with ROCK inhibitors completely abolished the increased cell plasma membrane strength and the Young’s modulus factors measured by AFM after inhibitors treatment reached similarly low levels in *shMAGI1* and *shLuc* controls (Supplementary Fig. S6C).

**Figure 4.**
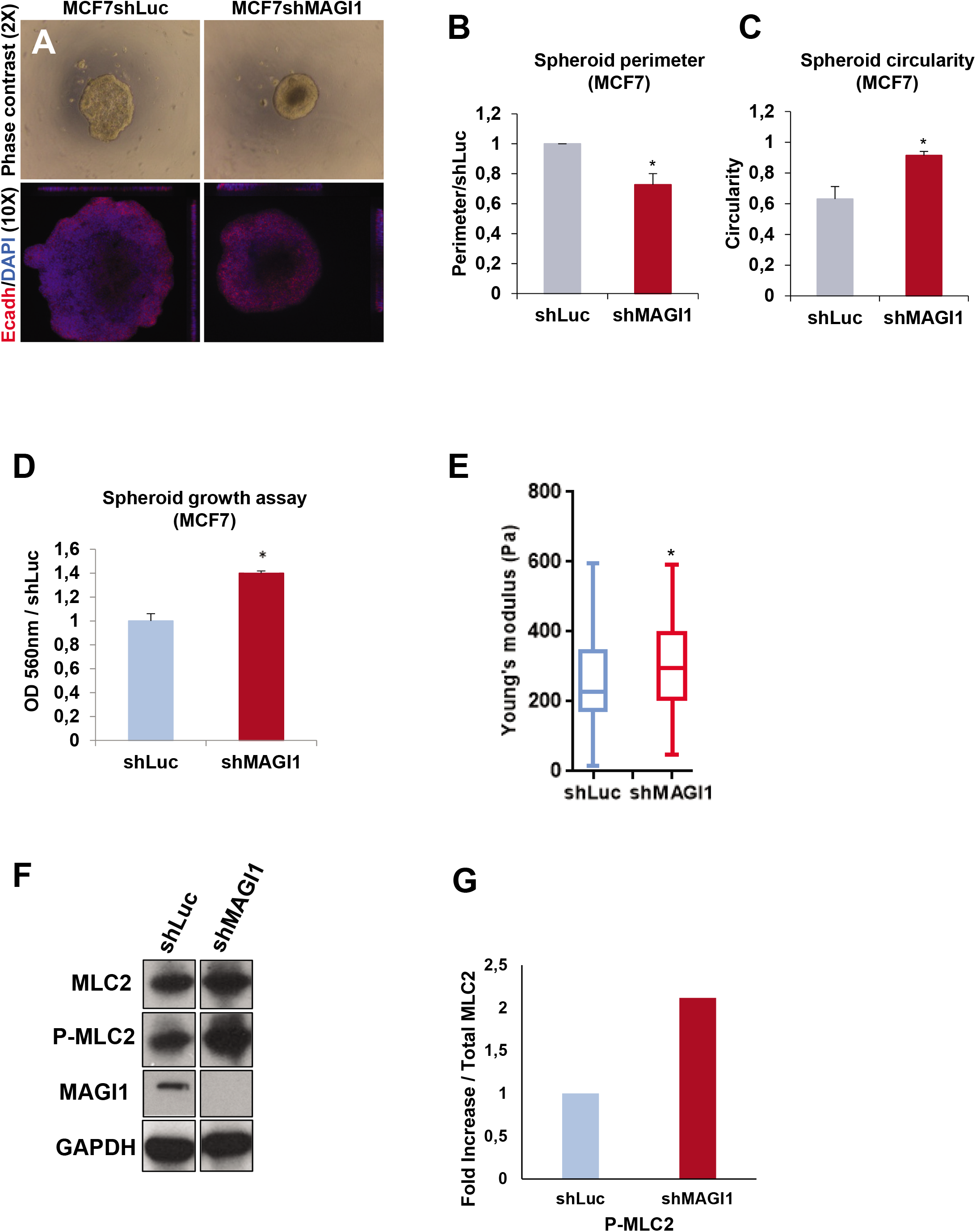
The loss of MAGI1 affects MCF7 cell compaction, ROCK activity, and compressive forces. (A) Representative phase contrast and fluorescence images of MCF7*shLuc* and MCF7*shMAGI1* cells grown in 3D spheroid cultures and stained with E-cadherin (red) and DAPI (blue). (B) Calculated perimeters for 3D spheroid cultures of MCF7*shMAGI1* normalized by the perimeter of MCF7*shLuc* cells. Bars represent mean ± SD (n=10 spheroids) of five independent experiments. Unpaired two-tailed Student’s t-test; * p < 0.05. (C) Calculated circularity for 3D spheroid cultures of MCF7*shLuc* and MCF7*shMAGI1* (calculations were done with the ImageJ software where a value of 1 is considered as a perfect circle). Bars represent mean ± SD (n=10 spheroids) of five independent experiments. Unpaired two-tailed Student’s t-test; * p < 0.05. (D) MTT assay (OD 560 nm) representing 3D spheroid cell growth of MCF7*shMAGI1* compared to MCF7*shLuc* cells 5 days after cell seeding. Bars represent mean ±SD (n=10 spheroids) of five independent experiments. Unpaired two-tailed Student’s t-test; * p < 0.05. (E) Elastic Young’s modulus (EYM) of cells: Hertz contact mechanics model for spherical indenters was used. In MCF7*shMAGI1* cells, the apical surface EYM is significantly elevated when compared to control MCF7*shLuc* cells. Data are represented as mean +/- SD: MCF7*shLuc* EYM= 258.8 +/-101, MCF7*shMAGI1* EYM= 302.4 +/- 91. Unpaired two-tailed Student’s t-test; * p < 0.05 (n=132 and 94 for MCF7*shLuc* and MCF7*shMAGI1* respectively). (F) Western blot analysis on whole protein extracts (n=3) of ROCK-specific ser19 phosphorylation of Myosin Light Chain 2, total MLC2 and MAGI1 in MCF7*shMAGI1* cell lines compared to MCF7*shLuc*. GAPDH was used as a loading control. Uncropped blots can be found in the Supplementary Information. (G) Quantification of the representative Western blot showing protein expression represented in panel E. Phosphorylated proteins were quantified as compared to their total protein counterparts to evaluate their activation.

### MAGI1 loss does not promote YAP nor β-catenin signaling

In epithelial cells, E-cadherin-based AJs regulate the activity of mechanosensitive pathways, and in particular the Hippo and Wnt/β-catenin pathways, two major pathways controlling cell proliferation and frequently mis-regulated during carcinogenesis ^8,46^. We thus explored whether these two pathways could be affected following MAGI1 depletion.

Monitoring Hippo pathway and YAP activity in stable MCF7*shMAGI1* showed that in the absence of *MAGI1*, total YAP and phosphorylated YAP (on both serine residues S127 and S397) accumulated in the cells while TAZ levels remained unchanged (Western blot analyses; Figure 5A). By immunofluorescence, we then showed that unlike what was observed for control cells grown at similar confluence, YAP did not accumulate in the nuclei of MCF7*shMAGI1* cells (Figure 5B). These findings were further confirmed in luminal BCa patients TMA, where we observed a strong correlation in ER+ tumors between MAGI1 apical membrane localization and YAP nuclear localization: only 27 % of the patients with no MAGI1 expression or mis-localized MAGI1 (no membrane staining) showed YAP nuclear localization while 92% of patients with membranous MAGI1 showed nuclear YAP (Figure 5C). Accordingly, *MAGI1* knockdown in MCF7 cells resulted in impaired YAP transcriptional activity with decreased expression of canonical YAP target genes such as *CTGF, CYR61, BIRC2, AXL*, and *AREG* (Figure 5D). The expression of *AMOTL2*, a YAP target gene encoding a negative YAP regulator, was increased after *MAGI1* knockdown, suggesting that despite its regulation by YAP, AMOTL2 transcription is also affected by other inputs in response to MAGI1 knockdown. Surprisingly, even though YAP phosphorylations on S127 and S397 are mediated by LATS kinases (Reviewed in ^47,48^), the loss of MAGI1 did not promote a higher activity of the Hippo pathway and the levels of phosphorylated activated LATS1 kinase and of phosphorylated MOB1 were unaffected (Figure 5A). This suggests that the elevated phosphorylated YAP were not the consequence of increased upstream Hippo pathway activity (MST kinases), but might reflect higher p-YAP stability, either because it was protected from destruction, or because it was not dephosphorylated as efficiently, or a combination of the two. These results are in agreement with earlier studies showing that E-cadherin junctional tension in epithelial cells promote YAP/TAZ nuclear exclusion either through α-catenin conformational change ^12–14^, or through Merlin/NF2-mediated cytoplasmic retention ^10^.

**Figure 5.**
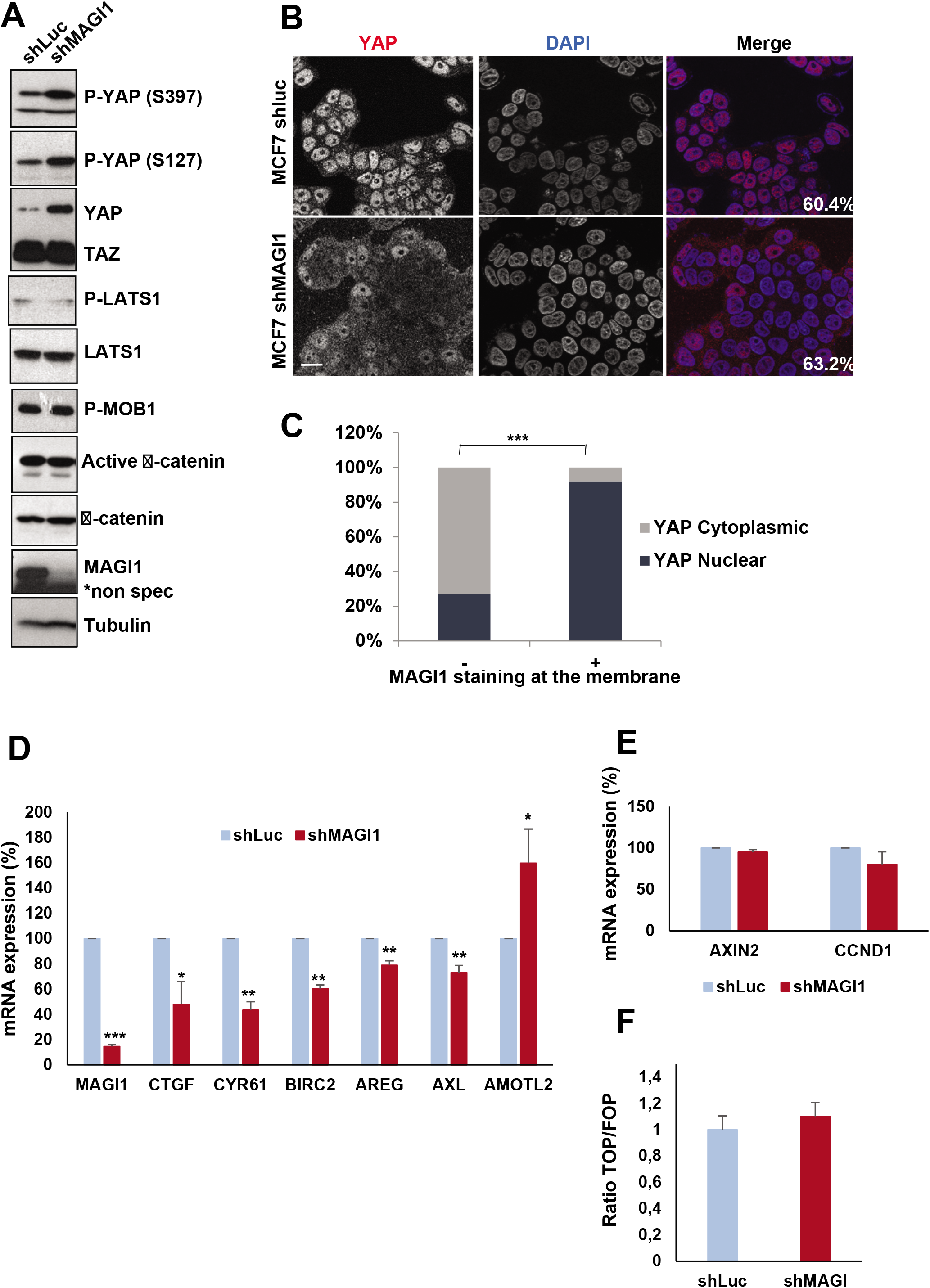
The loss of MAGI1 tumorigenic phenotypes are not associated with active YAP. (A) Western blot analysis on whole protein extracts (n=3) of phosphorylated YAP (S397 & S127), total YAP and TAZ, phosphorylated LATS1, total LATS1, phosphorylated MOB1, non-phosphorylated (active) β-catenin, total β-catenin and MAGI1 in MCF7*shMAGI1* cell lines compared to MCF7*shLuc*. Tubulin was used as a loading control. Uncropped blots can be found in the Supplementary Information. (B) Representative immunofluorescence images (n=3) of endogenous MAGI1 and YAP protein expression in MCF7*shLuc* and MCF7*shMAGI1* cells cultured to similar confluence (60.4 % and 63.2 % respectively). Scale bar=10 μm. (C) Quantification of MAGI1 and YAP staining in a tissue micro-array (TMA) from luminal breast cancer patients in a cohort of 28 patients, showing the strong correlation between MAGI1 membranous and YAP nuclear localizations; Fisher exact test, *** p<0.001. Histogram shows the percentages of patients having a nuclear or a cytoplasmic YAP staining according to the absence or the presence of MAGI1 membranous staining. (D) Real-time qRT-PCR showing the expression of canonical Hippo pathway target genes (CTGF, CYR61, BIRC2, AXL and AREG) as well as AMOTL2 in MCF7*shMAGI1* compared to the control MCF7*shLuc* cells and normalized with GAPDH expression. Data are presented as the means ± SD (n=3). Unpaired two-tailed Student’s t-test; * p < 0.05; ** p < 0.01; *** p < 0.001. (E) Real-time qRT-PCR showing the expression of two distinct Wnt pathway target genes (AXIN2 and CCND1) in MCF7*shMAGI1* compared to the control MCF7*shLuc* cells and normalized with GAPDH expression. Data are presented as the means ± SD (n=3). Unpaired two-tailed Student’s t-test revealed no statistical differences. (F) WNT reporter activity assay showing the TOP/FOP ratio in MCF7*shMAGI1* as compared to MCF7*shLuc* control cells. Data are presented as the means ± SD (n=3). Unpaired two-tailed Student’s t-test revealed no statistical differences.

Similarly, even though higher overall protein levels of β-catenin could be observed in MCF7*shMAGI1* cells compared to controls, this was not reflected by an increase in the active unphosphorylated β-catenin pool (Figure 5A). Accordingly, β-catenin transcriptional activity remained unchanged as evidenced by following by qPCR the expression of the AXIN2 and CCND1 canonical target genes or by TOP/FOP luciferase reporter assay (Figure 5E&F). These results are in agreement with studies showing that increased compression transduced by apical junctions in epithelial cells prevent β-catenin nuclear accumulation ^11,16^. Together, these results thus show that the loss of MAGI1 in luminal BCa cells does not promote YAP nor β-catenin signaling, and suggest that these pathways do not mediate the increased tumorigenicity of *MAGI1* deficient cells.

### p38 Stress Activated Protein Kinase mediates tumorigenicity of MAGI1 deficient cells

We thus asked which mechanisms mediate the increase in 2D cell proliferation and in anchorage-independent cell growth observed upon MAGI1 loss (see Fig. 1), focusing on other known oncogenic pathways. The ERK/MAP Kinase and JNK stress kinase pathways could be influenced by junctions and the forces they sense ^15,17,18^. In response to MAGI1 loss, we did not detect any alterations of the main oncogenic pathways: ERK1/2, JNK and Akt pathways (Figure 6A&B) in contrast with recently published results on MAGI1 function ^45^. Interestingly, the p38 stress kinase was strongly activated in MCF7*shMAGI1* compared to MCF7*shLuc*-: the amount of phosphorylated p38, normalized to total p38 protein level, was increased 2 fold (Figure 6A&B) leading to increased p38 activity as evidenced by the elevated expression of the p38 transcriptional target genes^49^ ATF3, SOX9, and GATA6 (SOX2 levels remained unchanged) (Figure 6C). Importantly, similar results regarding the activity of the different signaling pathways upon MAGI1 knock-down, and in particular the increased activity of p38, were obtained in other cell lines (T47D luminal BCa and colon HCT116, Supplemental Fig. S3G), showing that the effects observed are not restricted to MCF7 cells and represent a more general cellular response to MAGI1 loss. The p38 pathway is activated in response to a wide variety of cellular stresses and has been implicated either as a tumor-suppressor or as an oncogene in various cancers, including BCa ^50–54^. Using p38 specific inhibitors and siRNA mediated silencing, we then tested whether the p38 pathway could mediate, at least in part, the increased tumorigenicity of MCF7 luminal cells. In 2D cell growth and in soft agar anchorage-independent cell growth, MCF7*shMAGI1* treated with LY2228820 p38α/β inhibitor (a.k.a. Ralimetinib) grew less than untreated cells (2 fold suppression at 8 days of growth, and 2.5 fold decrease respectively; Figure 6D&E). Similar effects were obtained using another independent p38 inhibitor, ARRY-614 (p38 and Tie2 inhibitor), even though we observed higher toxicity (data not shown) and importantly using p38α knockdown by siRNA (Figure 6F&G), demonstrating that the effects observed were the consequence of p38 inhibition and not any unspecific action of the drugs. It should be noted that p38 inhibition also affected MCF7*shLuc* control cells (2D growth and soft agar), but the effect were far less pronounced than for MCF7*shMAGI1* cells, strongly supporting that the increased p38 activity is mediating, at least in part, the increased tumorigenicity upon MAGI1 depletion. Taken together, our results demonstrate the essential role of the p38 signaling pathway for MAGI1-deficient MCF7 luminal BCa cells tumorigenicity, consistent with previous report identifying the critical role of p38α in mammary tumorigenesis using mouse BCa models ^50^.

**Figure 6.**
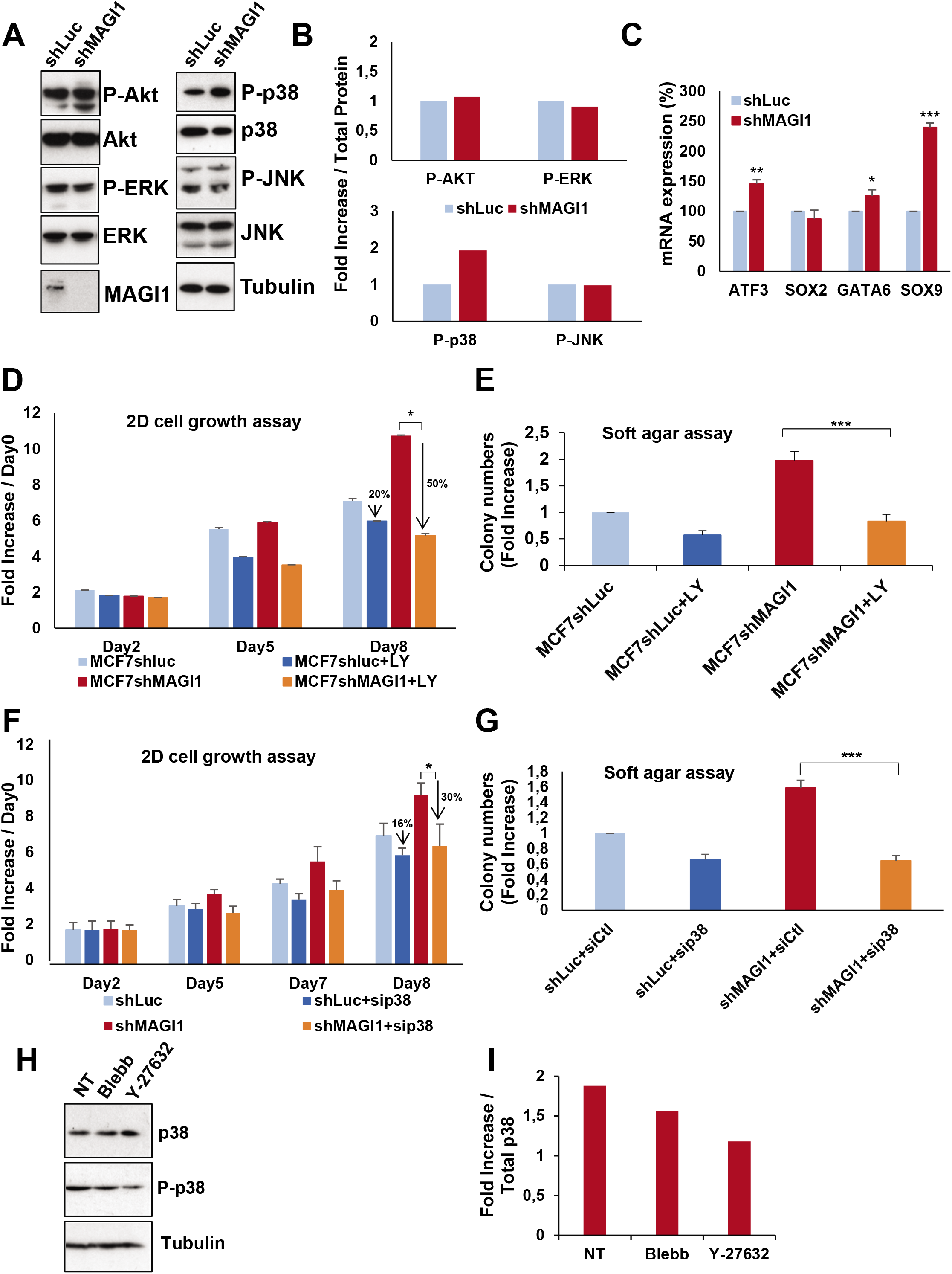
The loss of MAGI1 tumorigenic phenotypes are associated with p38 stress signaling pathway activation. (A) Western blot analysis showing protein expression and/or activation of Akt and MAPK proliferation signaling pathways as well as p38 and JNK stress signaling pathways in MCF7*shMAGI1 compared* to MCF7*shLuc* cells. Tubulin was used as a loading control. Western blot experiments have been repeated at least three times. Uncropped blots can be found in the Supplementary Information. (B) Quantification of the representative Western blot showing protein expression represented in panel A. Phosphorylated proteins were quantified as compared to their total protein counterparts to evaluate their activation. (C) Real-time qRT-PCR showing the expression of some p38 pathway target genes (ATF3, SOX2, SOX9 and GATA6) in MCF7*shMAGI1* compared to the control MCF7*shLuc* cells and normalized with GAPDH expression. Data are presented as the means ± SD (n=3). Unpaired two-tailed Student’s t-test; * p < 0.05; ** p < 0.01; *** p < 0.001. (D) 2D cell growth assay of MCF7*shMAGI1* treated, or not, with the p38 MAPK inhibitor LY2228820 (LY in the figure) compared to MCF7*shLuc*. Bars represent mean ± SD (n=6 wells as replicates) of a representative experiment (n=3). Unpaired two-tailed Student’s t-test; * p<0.05. Note the increased percentage of drop for MCF7*shMAGI1* cells at day 8. (E) Quantification of colony numbers of MCF7*shMAGI1* cells non treated or treated with LY2228820 grown in anchorage independent conditions (soft agar assay) and represented as fold increase compared to MCF7*shLuc* cells non treated or treated with the p38 inhibitor (LY in the figure). Data are presented as the means ± SD (n=3). Unpaired two-tailed Student’s t-test; *** p < 0.001. (F) 2D cell growth assay (MTT) representing 2D cell growth of MCF7*shMAGI1* and MCF7*shLuc* after transfection of *p38a*siRNA or *Ctl*siRNA. Bars represent mean ± SD (n=10 wells as replicates) of a representative experiment (out of 3). Unpaired two-tailed Student’s t-test; *p < 0.05. (G) Quantification of colony numbers of MCF7*shMAGI1* and MCF7*shLuc* after transfection of *p38a* siRNA or *Ctl* siRNA, grown in anchorage independent conditions (soft agar assay) and represented as fold increase compared to MCF7*shLuc* cells transfected with *Ctl* siRNA. Data are presented as the means ± SD (n=3). Unpaired two-tailed Student’s t-test; *** p < 0.001. (H) Western blot analysis showing protein levels of phosphorylated p38 and total p38 in MCF7*shMAGI1* cells treated or non-treated (NT) with Blebbistatin (Blebb) or Y-27632. Both inhibitors were used at 10 μM during 2 h and Tubulin that was used as a loading control. Uncropped blots can be found in the Supplementary Information. (I) Quantification of the representative Western blot showing protein expression and represented in panel E. Phosphorylated p38 was quantified as compared to total p38 to evaluate its activation.

Strikingly, treating MCF7*shMAGI1* cells with the Rho Kinase (ROCK) inhibitors Y-27632 or Blebbistatin that prevent Myosin-mediated F-actin tension, abolished p38 activation (Figure 6H&I) and largely decreased elastic cell Young’s modulus (Supplementary Fig. S6C), showing that increased ROCK activity and actin strength represent critical events to activate p38 signaling. However, due to their cell toxicity, these ROCK inhibitors treatments could only be maintained for short periods, preventing us to assay formally, whether releasing actin tension in MAGI1 deficient cells could suppress their tumorigenicity as would be expected from their effects on p38.

### MAGI1 interacts with the angiomotin family members AMOT and AMOTL2 at the junctions

In order to better understand how the loss of MAGI1 and the associated alterations in ROCK activity and in E-cadherin-based junctions could lead to p38 stress-kinase activation, we decided to focus our studies on the junctional MAGI1 molecular complexes. Several studies, including proteomic approaches, have described a physical association between MAGI1 and AMOT or AMOTL2, two members of the angiomotin family of apical scaffolds, acting both as junctions/actin linkers as well as negative regulators of YAP transcriptional activity ^41–43,55,56^. We thus, investigated whether AMOTs could mediate the effects of MAGI1 to restrict tumorigenesis in breast luminal cells.

First, using co-immunoprecipitation experiments, we confirmed that overexpressed Flag-tagged MAGI1 or endogenous MAGI1 interacted with endogenous AMOT and AMOTL2 in MCF7 luminal BCa cells (Figure 7A&B). Building on the availability of numerous constructs generated by the group of KL. Guan ^57^, we have then used AMOT as a paradigm to study the interaction between MAGI1 and AMOTs through co-immunoprecipitation experiments between Flag-tagged MAGI1 and HA-tagged AMOT constructs. Using point mutations affecting key structural Proline residues in the first and second WW motifs of MAGI1, we showed that the second WW motif was required for the interaction with AMOT, since co-immunoprecipitation was abolished by a point mutation on Proline residue 390 (mutated to Alanine MAGI1 P390A; Figure 7C&D). The first WW domain appeared dispensable as no change in binding was observed when mutating Proline 331 (MAGI1 P331A). Similar results were obtained performing co-immunoprecipitations on endogenous AMOTL2 (Figure 7E), showing that as for AMOT, the second WW domain of MAGI1 was required for AMOTL2 binding, confirming earlier findings ^43^. The N-terminus part of AMOT contains two PPXY motifs, which represent canonical WW domains interactors. Using HA-tagged AMOT point mutants, we then determined that the interaction between MAGI1 and AMOT occurred through the second PPXY motif (Figure 7F&G) since co-immunoprecipitation was lost in AMOT Y287A (Tyrosine 287 mutated to Alanine; the mutation AMOT Y242A did not show any effect). Together these results confirm using precise point mutations that MAGI1 binds to AMOT and AMOTL2 through its second WW domain, and at the level of the conserved second PPXY motifs of the N-terminal domains of AMOT family scaffolds.

**Figure 7.**
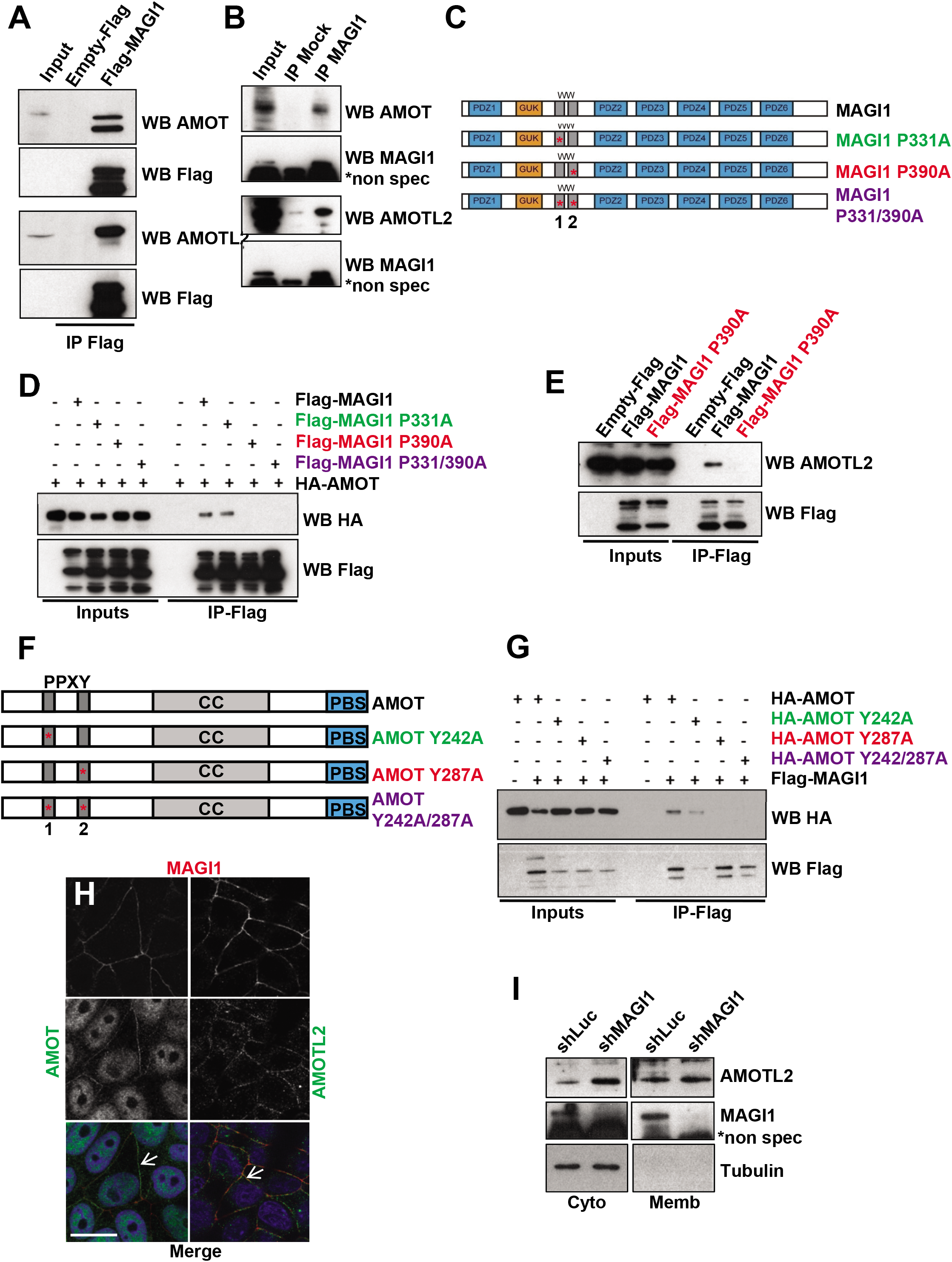
MAGI1 interacts with AMOT and AMOTL2. (A) Western blot of Flag-MAGI1 immunoprecipitates: Flag-MAGI1 transfections were performed in MCF7 cells and endogenous AMOT (upper panels) and AMOTL2 (lower panels) were revealed; Flag blotting was used as the immunoprecipitation control. Note that we could hardly detect Flag-MAGI1 by Western blot in the inputs, even though MAGI1 was well immunoprecipitated, due to sensitivity issues associated with single Flag tag in Flag-MAGI1 used in this typical experiment. Uncropped blots can be found in the Supplementary Information. (B) Western blot analysis of endogenous MAGI1 immunoprecipitates revealing the presence of endogenous AMOT (upper panels) and AMOTL2 (lower panels). MAGI1 blotting was used as the immunoprecipitation control (*shows non-specific bands with the anti-MAGI1 antibody). Uncropped blots can be found in the Supplementary Information. (C) Schematic of full length MAGI1 and of the point MAGI1 WW domain mutants used (P331A & P390A). (D) Western blot analysis of Flag-MAGI1 immunoprecipitates on protein extracts from MCF7 cells transfected with HA-AMOT and different Flag tagged MAGI1 constructs, showing that the interaction with AMOT occurred through the second WW domain of MAGI1 (interaction lost in P390A; red). Uncropped blots can be found in the Supplementary Information. (E) Western blot analysis of Flag-MAGI1 immunoprecipitates on protein extracts from MCF7 cells transfected with Flag-MAGI1 or Flag-MAGI1 P390A mutant, revealing the interaction of endogenous AMOTL2 with Flag-MAGI1 but not Flag-MAGI1 P390A (red). Uncropped blots can be found in the Supplementary Information. (F) Schematic of full-length AMOT and of the point AMOT PPXY mutants used (Y242A & Y287A). (G) Western-blot analysis of Flag-MAGI1 immunoprecipitates on protein extracts from MCF7 cells transfected with Flag-MAGI1 and different HA-AMOT constructs, showing that the interaction occurs through the second PPXY domain of AMOT (interaction lost in Y287A; red). Similar levels of interaction were obtained for HA-AMOT/Flag-MAGI1 and HA-AMOT Y242A/Flag-MAGI1 complexes (quantified using the ImageJ software). Uncropped blots can be found in the Supplementary Information. (H) Representative immunofluorescence images of endogenous staining for MAGI1 (red; top panels) co-localizing at the plasma membrane (white arrows) with AMOT (green; left) or AMOTL2 (green; right) in MCF7 cells. Note the non-specific nuclear staining for AMOT. Scale bar=10 μm. (I) Western blot analysis after subcellular fractionation showing the relative amount of AMOTL2 protein in the cytoplasmic and membrane fractions in MCF7*shMAGI1* compared to MCF7*shLuc* cells. Tubulin was used as a cytoplasmic control for the fractionation and for normalization. Uncropped blots can be found in the Supplementary Information.

Then, using immunofluorescence on MCF7 fixed cells, MAGI1, AMOT, and AMOTL2 localized in overlapping membrane domains corresponding to cellular junctions (Figure 7H; the AMOT nuclear staining is non-specific). Even though, MAGI1 colocalized with AMOTL2 and AMOT (Figure 7H, arrows), it was not required for their proper membrane localizations as they were unaffected in *shMAGI1*, suggesting that even though they physically interact once in proximity, they must reach their membrane localization independently (Figure 7I). Nevertheless, even though MAGI1 was not required for AMOTL2 localization, it did however control AMOTL2 levels, and AMOTL2 accumulated 1.8 fold in MCF7*shMAGI1* compared to controls as measured by western blots on whole protein extracts (Figure 3B). These results suggest that MAGI1 regulates the stability and/or degradation of AMOTL2, but using classic drugging approaches, we were unable to identify the mechanisms involved. It is noteworthy here that AMOTL2 mRNA levels were slightly increased in MAGI1-deficient cells (Fig. 5D), suggesting that the accumulation of AMOTL2 protein could also be a consequence of increased transcription.

Together our results confirm the physical interaction between MAGI1 and AMOTs and show that MAGI1 restricts the total AMOTL2 protein levels, through a mechanism that remains to be established.

### AMOTL2 mediates the increased tumorigenicity of MAGI1-deficient luminal BCa cells

We observed that in luminal BCa cells, the loss of MAGI1 triggered both i) an accumulation of AJs material including its binding partner AMOTL2, and ii) an activation of the p38 pathway driving an increase in tumorigenicity. We thus studied next whether these two different aspects were linked or independent consequences of MAGI1 loss. Generating double knockdown MCF7 cells for *MAGI1* and *AMOTL2*, we could show that the increased 2D growth and anchorage-independent growth after MAGI1 loss were suppressed by the knockdown of *AMOTL2* (Figure 8A&B). Consistently, *AMOTL2* knockdown abolished the accumulation of E-cadherin and the activation of the p38 stress kinase (phospho-p38 levels; Figure 8C&D). These results identify AMOTL2 accumulation as a critical mediator in the p38 activation and tumorigenesis induced by impaired *MAGI1*.

**Figure 8.**
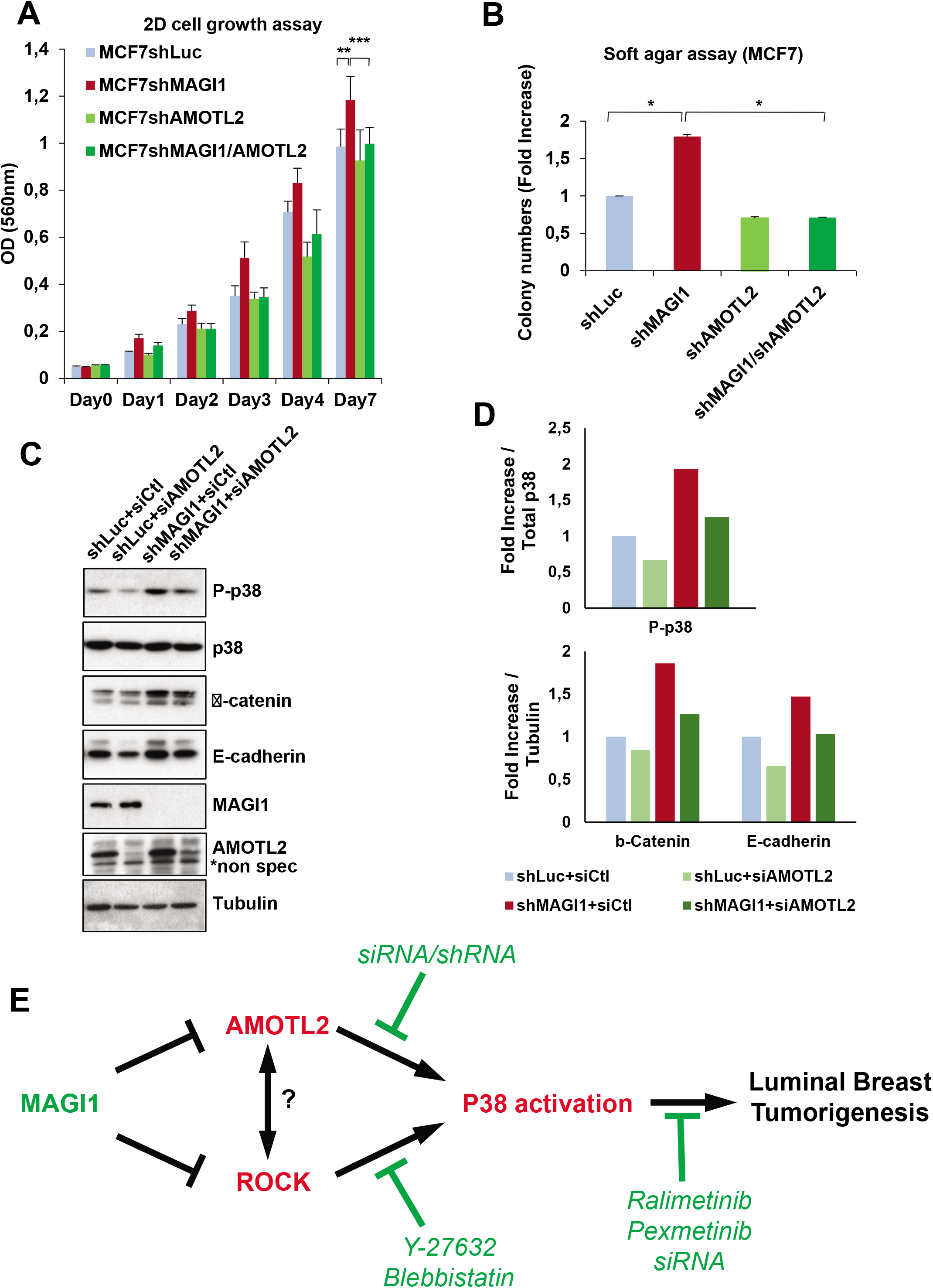
AMOTL2 mediates the effects of MAGI1 on junctions and on p38 signaling. (A) MTT assay (OD 560 nm) representing 2D cell growth of MCF7*shMAGI, MCF7shAMOTL2* and MCF7*shMAGI1/shAMOTL2* cells as compared to MCF7*shLuc*. Bars represent mean ± SD (n=10 wells as replicates) of a representative experiment (n=3). Unpaired two-tailed Student’s t-test; ** p<0.01; *** p < 0.001. (B) Quantification of colony numbers of MCF7 cells (*shMAGI1, shAMOTL2 & shMAGI1/shAMOTL2*) grown in anchorage independent conditions (soft agar assay) and represented as fold increase compared to MCF7*shLuc* cells. Data are presented as the means ± SD (n=3). Unpaired two-tailed Student’s t-test; *p < 0.05. Western blot analyses confirmed the 90 % knockdown for MAGI1 in MCF7*shMAGI1* and MCF7*shMAGI1/shAMOTL2* and QPCR data confirmed the 50 to 60 % knockdown for AMOTL2 in MCF7*shAMOTL2* and MCF7*shMAGI1/shAMOTL2* respectively (determined by RT-qPCR and/or Western Blot; Data not shown). (C) Western blot analysis showing protein expression and/or phosphorylation (activation) of p38 and JNK stress signaling pathways as well as junctions’ components in MCF7*shMAGI1* as compared to MCF7*shLuc* cells when *AMOTL2*siRNA was transfected. Tubulin was used as a loading control. Western blot experiments have been repeated at least three times. Uncropped blots can be found in the Supplementary Information. (D) Quantification of the representative Western blot showing protein expression and represented in panel C. Phosphorylated proteins were quantified as compared to their total protein counterparts to evaluate their activation and junctions’ proteins were quantified as compared to tubulin that was used as a loading control. Upper: Phosphorylated p38 / Total p38 protein quantification and Lower: protein / Tubulin quantification. (E) Model for the role of MAGI1 during Luminal BCa. MAGI1 prevents the accumulation of junctional AMOTL2 and E-cadherin as well as ROCK activity thus releasing cellular stiffness. Increased AMOTL2 and ROCK then activate p38 stress signaling responsible for the increased tumorigenicity of MAGI1-deficient cells. Anti and Pro tumorigenic events are highlighted in green and red respectively.

Together, these results show that both AMOTL2 and ROCK activity are required for p38 activation in MAGI1 deficient luminal BCa cells. They support a model in which, the AMOTL2 accumulation in MAGI1 deficient cells governs an increase of E-cadherin-based junctions, which together with ROCK activity and cortical actin tension, activates the tumorigenic activity of the p38 signaling pathway (Figure 8E).

## DISCUSSION

In this study, we explored the function of the junctional component MAGI1 in breast carcinogenesis and showed that the loss of MAGI1 in luminal BCa cell lines promoted tumorigenesis. At the cellular level, we showed that the loss of MAGI1 induced an accumulation of AJ components and increased cell compression behaviors. Importantly, we showed that the increased tumorigenicity of MAGI1 deficient luminal BCa cells was mediated by p38 stress signaling. Molecularly, we confirmed that MAGI1 interacted with the AMOT family of apical scaffolds and actin linkers, and showed that decreasing *AMOTL2* levels, completely suppressed the effect of *MAGI1* loss on the activation of p38 and on tumorigenicity. Finally, we provided evidence that inhibiting ROCK activity could alleviate the p38 activation, supporting a model in which MAGI1 suppresses luminal BCa by preventing both ROCK and AMOTL2-mediated Junction dysfunction, and subsequent p38 stress signaling.

The loss of MAGI1 led to an increase in tumorigenicity, and an accumulation of AJ material, in particular E-cadherin, reminiscent to the role we described previously for the unique MAGI homologue in *Drosophila* ^24^. This correlation could appear counterintuitive as E-cadherin knockdown promotes invasion and migration in cultured cells ^58^, and is critical during EMT and for aggressiveness (recently reviewed in ^59^). However, in several cancers including some BCa, E-cadherin expression is maintained with a proposed tumor supporting role ^60,61^ and recent mouse models of luminal and basal invasive ductal carcinomas have demonstrated that mammary cancer cells proliferation, resistance to apoptosis, and distant metastatic seeding potential, require E-cadherin ^62^. These observations reconcile the high E-cadherin levels and increased tumorigenicity observed upon *MAGI1* loss, and suggest that at least in specific phases of tumorigenesis (e.g. mass proliferation, colony formation), E-cadherin could play a positive role during BCa.

One striking feature of MAGI1 depleted cells is the protein accumulation of AMOTL2. Even though the exact mechanisms remain to be described, the stability of the related AMOT is controlled by the activity of the RNF146/Tankyrase pathway ^63,64^, and by phosphorylation by the Hippo pathway core kinase LATS1 (reviewed in ^37^). Importantly AMOTL2, a binding partner of MAGI1, is required for the increased tumorigenicity of MAGI1-depleted luminal BCa cells, showing that AMOTL2 is a critical mediator in this context. AMOTs are apical scaffolds playing important roles in cadherin-based junction regulations and its linkage to the actin cytoskeleton ^26,35,36^. Previous reports have shown that AMOTL1 is overexpressed in invasive ductal carcinomas, leading to invasive behaviors. However, this effect of AMOTL1 appears specific to ER-negative subtypes ^65^. AMOTL1 is not expressed in breast luminal lineages, in which AMOTL2 is the most abundant (twice more abundant than AMOT in MCF7 cells; our unpublished observation). We propose thus that in ER-positive luminal breast lineages, AMOTL2 overexpression could play similar roles as marker and mediator of tumorigenesis. Interestingly, a short isoform of AMOTL2 (AMOTL2 p60 in contrast to the long AMOTL2 p100) was reported overexpressed in *in situ* and invasive ductal carcinoma cells ^66^. The AMOTL2 p60 isoform lacks the PPxY domain critical for its interaction with MAGI1, and it is possible this isoform is thus insensitive to MAGI1-dependent destabilization, even though this remains to be studied. Hypoxia and Fos signaling govern the specific expression of AMOTL2 p60 ^66^, and the accumulation of AMOTL2 in the context of *MAGI1* mutation could thus represent an alternative mechanism for AMOTL2 expression, extending the relevance of AMOTL2 up-regulation during carcinogenesis.

AMOTL2 couples adhesion complexes to the actin cytoskeleton to allow F-actin tension and thus morphogenesis in endothelial cells (coupling with VE-cadherin ^67^) and in epithelial cell during embryogenesis (E-cadherin ^56^). Importantly, the specific cellular morphology defects in AMOTL2 knockdown could be mimicked by inhibiting ROCK kinase and Myosin phosphorylation showing that AMOTL2 and actin tension are linked ^35^. Relaxing actin tension using two ROCK inhibitors (Y-27632 and Blebbistatin), or preventing AMOTL2 accumulation (siRNA), both robustly suppressed p38 activation, a key feature mediating tumorigenesis after MAGI1 loss. So how could AMOTL2 and ROCK interact to mediate the effects of *MAGI1* loss? They could either (i) be two independent consequences of MAGI1 loss or (ii) they could be linked causally. Due to the toxicity of long term exposures to ROCK inhibitors, we were unable to study whether AMOTL2 stability could be a consequence of ROCK activity. Preliminary results would indicate that AMOTL2 does not control ROCK activity (data not shown), suggesting that ROCK might act upstream, or in parallel to AMOTL2, but more experiments are needed to better determine how MAGI1 controls ROCK activity, and how ROCK and AMOTL2 regulate p38 activity.

p38 signaling is activated by many upstream signals including a wide variety of environmental stresses such as UV, genotoxic agents, heat shock, or hyperosmotic conditions ^18^. During osmotic shock, the cellular cortex is subjected to external pressure reminiscent to compressive forces, and we propose therefore that E-cadherin and AMOTL2 enrichment, together with increased ROCK activity and actin tension result in similar compressive forces that could represent a new stress signal activating p38. Indeed, amongst the genes regulated transcriptionally by p38, SOX9, a gene mostly responsive in osmotic shock^49^, was the most robustly activated upon MAGI1 knock-down. Amongst the four different p38 kinases (p38α,β,γ,δ), P38α is the most ubiquitously expressed, p38 signaling has been implicated either as a tumor suppressor or as an oncogene depending on cancer type and stage ^18,52–54^. In BCa, targeting p38δ resulted in a reduced rate of cell proliferation and enhanced cellmatrix adhesion ^68^. More importantly, BCa mouse models, demonstrated the critical role of p38α (Mapk14) during the initiation, and proliferation of mammary tumors. The tumor promoting role of p38α involves a protective role against deleterious DNA damage in the mammary epithelial cancer cells^50^. We observed in *MAGI1*-deficient cells an increased proportion of cells in S-phase, indicating that these cells might have a slower S-phase. Slow S-phase is classically observed when cells have to overcome a high level of DNA damage. These results highlight the need for further studies to better understand the link between MAGI1 loss, p38 activation, DNA damage, and breast tumorigenesis.

Besides elevated p38 signaling, MAGI1-deficient cells exhibited low YAP nuclear activity, consistent with earlier reports showing that in epithelial cells with high E-cadherin and/or under compressive forces α-Catenin and NF2/Merlin act to exclude YAP from the nucleus ^10^. Here we demonstrated the critical role of AMOTL2 downstream of MAGI1 to mediate E-cadherin accumulation. Since AMOTs have been shown to trap YAP in the cytoplasm ^57^, further studies are thus required to better understand the interactions between MAGI1/AMOTL2 and α-Cat/NF2 in the control of YAP localization and activity. This low YAP signaling could appear surprising since the oncogenic role of nuclear YAP is well established ^69^. However, the oncogenic role of YAP in luminal BCa remains debated. While elevated YAP/TAZ activity gene signatures have been reported to correlate with more aggressive BCa ^70–72^, aggressive BCa are enriched in basal/triple negative sub-types. Furthermore, several studies report conflicting or no correlation between YAP staining levels and clinical outcomes in BCa patients (reviewed in ^69^) suggesting that YAP/TAZ levels and nucleo/cytoplasmic distributions in luminal BCa could be re-examined.

## METHODS

### Plasmids, mutant constructs and shRNA cloning

All expression plasmids were generated with the Gateway system (Invitrogen). The Entry vector *pDONR223-MAGI1* was a gift from William Hahn (Addgene plasmid #23523) ^73^ and it was used to introduce the different point mutations in *MAGI1* by mutagenesis PCR using PfuTurbo (Agilent). All the Gateway destination vectors are listed in the supplementary methods. *pEZY-EGFP-MAGI1* was constructed by Gateway recombination between the *pDONR223-MAGI1* and the *pEZY-EGFP* destination vector that was a gift from Zu-Zhu Zhang (Addgene plasmid #18671) ^74^. *pcDNA3 HA-AMOT, pcDNA3 HA-AMOT Y242A, pcDNA3 HA-AMOT Y287A* and *pcDNA3 HA-AMOT Y242/287A* were gifts from Kunliang Guan (Addgene plasmids #32821, #32823, #32824 and #32822 respectively) ^38^.

shRNA directed against human *MAGI1* and *AMOTL2* were constructed and cloned in the pSIREN-RetroQ vector (TaKaRa) according to the manufacturer’s conditions between BamHI and EcoRI cloning sites as previously described for other genes ^75^. Targeted sequences are listed in the supplementary methods section.

### Cell culture and cell transfection

MCF7, T47D and HCT116 human cell lines originated from the *TumoroteK* bank (SIRIC Montpellier Cancer) and have been authenticated by STR (Short-tandem repeat) profiling and certified mycoplasm-free. Details for culture are added in the supplementary methods.

Plasmid transfections were performed in MCF7 cells using Lipofectamine2000 reagent (Invitrogen) according to the manufacturer’s directions. Cell extracts were assayed for protein expression 24-48 hours post-transfection. Concerning *AMOTL2*siRNA and *p38α*siRNA transfection (Dharmacon siGENOME #M-01323200-0005 and #L-003512-00-0005 respectively), a final concentration of 100 nM was used using Lipofectamine2000 reagent as well.

### Western blotting

Proteins issued from transfected MCF7, MCF7shRNA, T47DshRNA or HCT116shRNA cell lines were extracted, analyzed by SDS-PAGE. For dilutions and antibodies references, please refer to the supplementary methods.

### RNA extraction, reverse transcription and real time RT-qPCR

RNAs were extracted from cells using RNeasy plus kit (Qiagen) and cDNAs were prepared starting from 1 μg of RNAs using the Superscript III reverse transcriptase (Invitrogen) and following manufacturer’s directions. Real time quantitative PCR was then performed on cDNAs using SYBR Green I Mastermix (Roche) according to the manufacturer’s conditions on a Light Cycler 480 device (Roche). All primers used are listed in the supplementary methods.

### Patients and TMA construction

Breast cancer samples were retrospectively selected from the Institut régional du Cancer de Montpellier (ICM) pathology database using the following inclusion criteria: chemotherapy-naïve at the time of surgery and estrogen (ER), progesterone (PR) receptor and HER2 status available. TMA construction are detailed in the supplementary methods section. Tumor samples were collected following French laws and declared to the French Ministry of Higher Education and Research (Biobanque BB-0033-00059; declaration number DC-2008–695). All patients were informed about the use of their tissue samples for biological research and the study was approved by the local translational research committee (CORT).

### Immunohistochemistry

Four μm thin sections of the TMA were mounted on Flex microscope slides (Dako) and allowed to dry overnight at room temperature before immunohistochemistry (IHC) processing. For further details, please refer to the supplementary methods.

### Immunofluorescence

Cells seeded on glass coverslips were fixed 10 min in paraformaldehyde (4 %), before being permeabilized in PBS / 0.1% TritonX-100 for 10 min. After blocking in PBS / 0.5% BSA, cells were incubated with primary antibodies overnight at 4C. Antibodies used are listed in the supplementary methods. Secondary Alexa Fluor Antibodies (1/200; Invitrogen) were used as described previously ^24^ for 1 hour at room temperature before

**Atomic** mounting the coverslips with Vectashield (Vector Laboratories #H-1200) and imaging on Zeiss Apotome or Leica Thunder microscopes.

### Force Microscopy (AFM) measurements

Cells were plated on glass 35mm fluorodish at intermediate (1/3) dilution density, grown overnight, and placed in 2% serum medium before AFM experiments. Cells grew as clusters of cells. AFM Force Spectroscopy (AFM-FS) experiments ^76^ on living cell clusters were performed on a JPK Nanowizard 4 microscope equipped with a CellHesion stage (JPK). The AFM is mounted on an inverted Axiovert 200M microscope system (Carl Zeiss) equipped with a 100x (1.5NA, plan-Apochromat) objective lens (Carl Zeiss). We employed a heating module (JPK) placed at the sample level to maintain cells at the physiological temperature of 37°C during measurements. We used CP-CONT-SIO-D cantilevers with 10μm colloidal beads as tip (NanoAndMore). Cantilever stiffness and optical lever sensitivities were both calibrated in liquid environment using the Contact-Free-Method provided by JPK AFM, and based on a mix of a thermal and Sader^77^ calibration methods. For calibration details, please refer to the supplementary methods. At least 5 force maps were acquired in each experiment on each cell category and 3 separated experiments were performed. Analyses were carried out using the JPK AFM data processing software. The elastic Young’s modulus (E; Pa) was evaluated by fitting each force versus tip-cell distance curve with the Hertz contact model for indenting an infinite isotropic elastic half space with a solid sphere as described in ^78^.

### Mammosphere culture

MCF7*shLuc* and MCF7*shMAGI1* cell suspensions were obtained by trypsinization and then filtered using 30μm cell strainer (Miltenyi). Filtered cells were immediately seeded in 96 well plates coated with 1% Poly (2-hydroxyethyl methacrylate; Sigma Aldrich) at a density of 200 or 50 cells per well. Cells were grown in MEBM basal medium supplemented with B27 without vitamin A (50X), 20 ng/mL EGF, 20 ng/mL bFGF and 4 μg/mL heparin. Cells were incubated at 37 °C with 5% CO_2_ during 5 days. For mammosphere acquisition and analysis, please refer to the supplementary methods.

### Immunoprecipitation and co-immunoprecipitation

Protein extracts were prepared in lysis buffer (NaCl 150 mM, Tris pH 7.4 10 mM, EDTA 1 mM, Triton X-100 1%, NP-40 0.5%, cOmplete, EDTA-free protease inhibitors (Roche #11873580001) for 30 min on ice before centrifugation. Immunoprecipitations were performed overnight at 4°C on a rocking wheel using either mouse anti-MAGI1 antibody for endogenous immunoprecipitations or anti-HA antibody and EZview Red anti-FLAG M2 affinity gel (Sigma-Aldrich #F1804) for co-immunoprecipitations. Protein G sepharose was then added to the MAGI1-or HA-immunoprecipitates for 1 hour at 4°C before extensive washes. Concerning the FLAG immunoprecipitation, washes were performed followed by protein elution by competition with 3XFLAG peptide (150 ng/μL final concentration) during 1 hour at 4°C. The different immunoprecipitates were then submitted to Western blotting for detection of protein complexes.

### Subcellular fractionation - Isolation of cytoplasmic, nuclear and membrane protein fractions

Subcellular fractionation of cultured human cell lines was performed as previously described on-line (http://www.bio-protocol.org/e754).

### Cell growth assay

2D cell growth assay was analyzed using MTT (Thiazolyl Blue Tetrazolium Bromide (Sigma Aldrich #M2128)). Briefly, cells seeded in 96 well plates were stopped from day 0 to day 7 by incubation with MTT solution (0.5 mg/mL final concentration) during 4 hours at 37 °C. Before reading OD at 570 nm, cells were incubated with DMSO to solubilize the formazan crystals formed in the presence of MTT.

### 3D Spheroid formation and measures

10 000 cells were seeded in Ultra-Low Attachment 96 well plates (Costar) cultured in their regular medium supplemented with 15 % FCS. After 3 days, all cells formed 3D spheroids that can be further analyzed by perimeter and circularity measurements. Pictures of spheroids were taken 3 days after seeding cells to assay the ability of cell lines to form spheroids and both the perimeters and the circularity of spheroids were calculated. Circularity was calculated (4px(area/(perimeter)^2^) where 1 is considered as a perfect circle.

### p38 MAPK inhibitor treatment

The different MCF7*shRNA* cell lines were plated at 3×10^6^ cells per 60mm plates. The day after, 10 μM of Ralimetinib (LY2228020; Selleckchem #S1494) was added and 24 h later cells were collected and counted to be plated for the different experiments (Ralimetinib treatment is maintained throughout the experiments). For Soft agar assay, 10 000 cells were plated in 12-well plates before proceeding to the assay detailed below. The same experiments were conducted with another p38 inhibitor using 10 μM of Pexmetinib (ARRY-614; Selleckchem #S7799). SRB proliferation assay was performed as detailed in the supplementary methods.

### ROCK inhibitors treatment

The treatments of MCF7*shRNA* cell lines with either Blebbistatin (Selleckchem #7099) or Y-27632 (Selleckchem #S1049) were both performed at a final concentration of 10 μM and cells were treated during 2 hours before being lysed for Western blot analyses.

### Soft Agar assay

To assess anchorage independent growth, soft agar assays were performed. A first layer of 0.8% Noble agar (Sigma Aldrich #A5431) was prepared in 12-well plates and left at 4C during 1 hour for the agar to solidify gently. The plate was kept at 37C before the cell-containing agar layer was prepared. 5,000 or 10,000 cells were imbedded in the second layer containing 0.4% Noble agar on top of the first layer. When the second layer was solidified, fresh medium was added on top of the cells. MCF7 cell lines were cultured during 14 to 21 days before Crystal violet coloration (0.01% final concentration; Sigma Aldrich #C0775) or MTT assay (1 mg/mL final concentration).

### Primary tumor growth assay

MCF7shRNA and HCT116shRNA cell lines were injected subcutaneously in six-week-old female athymic Nude-Foxn1 mice (Envigo). Both flanks of each mouse were injected with cells mixed with Matrigel (10^6^ cells injected for HCT116; 10×10^6^ cells injected for MCF7). Mice injected with MCF7 cells were subjected to neck dropping with 50μl of β-estradiol (1.5 mg/mL diluted in ethanol) to stimulate MCF7 cell growth. Animal procedures were performed according to protocols approved by the French national committee on animal care and this study was carried out in compliance with the ARRIVE guidelines.

## ACKNOWLEDGEMENTS

We acknowledge the contribution of G. Froment, D. Nègre and C. Costa from lentivectors’ production facility of SFR Biosciences (UMS3444/CNRS, US8/Inserm, ENS Lyon, UCBL, France). We thank the ABIC (IRCM), MRI (IRCM), MGC (IGMM) platforms and M. Larroque from the URT (IRCM, Montpellier, France). We thank V. Chambon and T. Soirat for technical help with the mammosphere formation assay and PE Milhiet (CBS, Montpellier, France) for helpful discussion and AFM set up. All plasmids obtained from Addgene are detailed in the Material and Methods section and all references have been included. We also thank Drs C. Gongora, L. Holmgren, N. Tapon, and B. Zhao for discussion, tools and advice. This work was supported by grants from Fondation ARC and Ligue Régionale Contre le Cancer (34) to LHM and AD. DK is supported by Ligue Nationale Contre le Cancer. VS and EB were supported by Fondation de France. MM was supported by Fondation ARC.

## AUTHOR CONTRIBUTIONS

DK, EBM, MM, VS, YBS, LC, EF, FBM, BO, CB and LHM performed experiments. LHM and AD designed the experiments. DK, EBM, MM, JC, BO, LHM, and AD analyzed the data. DK, PL, CG, JC, CT, AM, LHM, and AD interpreted the data. LHM and AD wrote the manuscript.

