## Supplemental Material for "MAGI1 inhibits the AMOTL2/p38 stress pathway and prevents luminal breast tumorigenesis"

[1] IRCM, Univ Montpellier, Inserm, ICM, Montpellier, France. [2] Translational Research Unit, ICM, Montpellier, France. [3] CBS, Univ Montpellier, CNRS, Inserm, Montpellier, France.

[+] Co-second authors: E.B.M. and M.M.

[&] Co-last authors and co-corresponding authors: L.H.M. and A.D.

* Lead contact: A.D.

#### SUPPLEMENTARY FIGURE LEGENDS

##### Supplementary Figure S1

(A) Western blot showing MAGI1 expression in different breast cancer cell lines (T47D and MCF7 from the luminal A sub-type, MDA-MB-468 and BT549 from basal sub-type and ZR75 from luminal B sub-type. Tubulin was used as loading control.

(B) Kaplan-Meier survival curves and log-rank analysis. Relapse-free survival (RFS) curves for 841 ER positive breast cancer patients as a function of MAGI1 expression. Patients were split according to median expression. Data for MAGI1 (225474_at) were obtained using the KM plotter website at http://kmplot.com.

(C) Representative immunohistochemical staining of MAGI1 performed in normal human breast samples. Brown staining indicates positive immunoreactivity and arrows show MAGI1 staining.

(D) Western blot analysis of MAGI1, MAGI2 and MAGI3 expression in MCF7*shMAGI1(1-1)* and MCF7*shMAGI1(3-1)* showing respectively 50% and 100% knockdown as compared to MCF7*shLuc* control cell line. Tubulin was used as a loading control.

(E) Representative immunofluorescence images showing MAGI1 staining (red) in the *MCF7shMAGI1 (1-1)* and *(3-1)* and in MCF7*shLu*c cell lines. Scale bar=10 µm.

(F) Real-time qRT-PCR showing the expression of *MAGI1*, *MAGI2* and *MAGI3* mRNA in MCF7*shMAGI1(1-1)* and in MCF7*shMAGI1(3-1)* compared to the control MCF7*shLuc* cell line. Data are presented as the means ± SD. Three independent experiments; unpaired two-tailed Student's t-test (* $p < 0.05$; n.s. not significant).

##### Supplementary Figure S2

(A) MTT assay (OD 560 nm) representing 2D cell growth of MCF7*shMAGI1(3-1)* and MCF7*shMAGI1(1-1)* as compared to MCF7*shLuc* cells. Bars represent mean ± SD; n=10 wells as replicates) of a representative experiment (out of 4). Unpaired two-tailed Student's t-test; * $p<0.05$; ** $p < 0.01$; *** $p < 0.001$.

(B) Quantification of colony numbers of MCF7*shMAGI1(3-1)* and MCF7*shMAGI1(1-1)* cells grown in anchorage independent conditions (soft agar assay) and represented as fold increase compared to

MCF7*shLuc* cells. Data are presented as the means ± SD (n=3). Unpaired two-tailed Student's t-test; ** $p < 0.01$.

(C) Representative phase contrast images of MCF7*shLuc,* MCF7*shMAGI1(3-1)* and MCF7*shMAGI1(1-1)* cells grown in 3D spheroid cultures.

(D) Calculated circularity for 3D spheroid cultures of MCF7*shLuc,* MCF7*shMAGI1(3-1)* and MCF7*shMAGI1(1-1)* (calculations were done with the ImageJ software where a value of 1 is considered as a perfect circle). Bars represent mean ± SD (n=10 spheroids) of five independent experiments. Unpaired two-tailed Student's t-test; * $p < 0.05$.

(E) Real-time qRT-PCR showing the expression of MAGI1, CTGF, CYR61, BIRC2, AREG and AMOTL2 mRNA in MCF7*shMAGI1(1-1)* and in MCF7*shMAGI1(3-1)* compared to the control MCF7*shLuc* cell line. Data are presented as the means ± SD. Three independent experiments; unpaired two-tailed Student's t-test (* $p < 0.05$; ** $p < 0.01$; *** $p < 0.001$).

(F) Western blot analysis on whole protein extracts (n=3) analyzing the expression of junctional proteins, proteins implicated in EMT, Hippo, Wnt, p38 and ROCK signaling pathways in MCF7*shMAGI1(1-1)* and in MCF7*shMAGI1(3-1)* compared to control MCF7*shLuc* cells. Tubulin was used as a loading control.

**Supplementary Figure S3**

(A) Phase contrast images showing wound healing assay of MCF7*shMAGI1* compared to MCF7*shLuc* cells. These are representative images of four different experiments.

(B) Quantification of the wound area for MCF7*shMAGI1* represented as fold increase compared to MCF7*shLuc*. Data are presented as the means (n=4) ± SD. Unpaired two-tailed Student's t-test revealed no statistical differences.

(C) Migration assay of MCF7*shMAGI1* cells performed in Boyden chambers and represented as the means (n=3) ± SD of migrating cells (%) compared toMCF7*shLuc* control cells. Unpaired two-tailed Student's t-test revealed no statistical differences.

**Supplementary Figure S4**

(A) MTT assay (OD 560 nm) representing 2D cell growth of T47D*shMAGI1* as compared to T47D*shLuc* cells. Bars represent mean ± SD (n=10 wells as replicates) of a representative experiment (n=3), showing similar increased cell growth as observed for MCF7 cells. Unpaired two-tailed Student's t-test; *** p < 0.001.

(B) Upper: quantification of the number of colonies of T47D*shMAGI1* grown in anchorage independent conditions (soft agar assay) represented as fold increase compared to T47D*shLuc* cells. Data are presented as the means (n=3) ± SD. Unpaired two-tailed Student's t-test; ** p < 0.01. Below: western blot analysis on whole protein extracts from cells used on the left showing the efficient MAGI1 protein knockdown; Tubulin was used as a loading control.

(C) Calculated perimeters for 3D spheroids cultures of T47D*shMAGI1* normalized by the perimeter of T47D*shLuc* cells. Bars represent mean ± SD (n=10 spheroids) of three independent experiments. Unpaired two-tailed Student's t-test; * p < 0.05.

(D) Calculated circularity for 3D spheroid cultures of T47D*shLuc* and T47D*shMAGI1*. Calculations were done with the ImageJ software where a value of 1 is considered as a perfect circle. Bars represent mean ± SD (n=10 spheroids) of three independent experiments. Unpaired two-tailed Student's t-test; * p < 0.05.

(E) Upper: quantification of the number of colonies of HCT116*shMAGI1* grown in anchorage independent conditions (soft agar assay) represented as fold increase compared to HCT116*shLuc*. Data are presented as the means (n=3) ± SD. Unpaired two-tailed Student's t-test; * p < 0.05. Below: analysis on whole protein extracts from cells used on the left showing the efficient MAGI1 protein knockdown; Tubulin was used as a loading control.

(F) Primary tumor growth of HCT116*shLuc* and HCT116*shMAGI1* cells injected subcutaneously in nude mice. Primary tumor growth was assessed by measuring the tumor volume over time until the tumors were too big and the mice had to be euthanized. Bars correspond to the average ± SEM (n=8 mice). Unpaired two-tailed Student's t-test (* p < 0.05).

(G) Western blot analyses on whole protein extracts (n=3) showing Akt, Wnt, p38 and ROCK signaling pathways along with AMOTL2 and E-cadherin total protein expression in T47D and HCT116 shRNA cell lines. GAPDH was used as a loading control.

**Supplementary Figure S5**

(A,D,G,J) Representative immunohistochemical staining of CD45 (~~staining~~ immune cells; A), CD31 (endothelial cells; D), $\alpha$-SMA (fibroblasts; G), or MAGI1 in primary tumors obtained after subcutaneous engraftment of either MCF7*shLuc* (left) or MCF7*shMAGI1* cells (right). Brown staining indicates positive immunoreactivity. Scale bar=250 µM (A,D,G) or 10 µm (J).

(B,C,E,F,H,I) Quantification of the immunohistochemical analyses in A,D,G and showing the percentage of primary tumor cells stained positively for CD45 (B), CD31 (E), or $\alpha$-SMA (H) and the normalized mean intensity of CD45 (C), CD31 (F), or $\alpha$-SMA (I) positive cells in primary tumors derived from MCF7*shMAGI1* compared to MCF7*shLuc* cells. Bars correspond to the mean ± SD (n=4 for shLuc and n=5 for shMAGI1). Unpaired two-tailed Student's t-test; n.s. not significant.

**Supplementary Figure S6**

(A) Representative immunofluorescence images of endogenous p-MLC (green staining) and actin (phalloidin red staining) in MCF7*shLuc* and MCF7*shMAGI1* . Scale bar=10 µm.

(B) Quantification of actin-associated p-MLC positive foci observed and counted in panel B (n=3). Bars represent mean ± SD (n=3). Unpaired two-tailed Student's t-test; * $p < 0.05$.

(C) Elastic Young's modulus (EYM) of cells treated, or not, with Y-27632 (10µM for 45 min): Hertz contact mechanics model for spherical indenters was used. In treated cells, the apical surface EYM is significantly decreased when compared to untreated cells. Data are represented as mean +/- SD : MCF7*shLuc* untreated EYM= 248.4 +/- 133.5, treated with Y-27632 EYM= 112.0 +/- 81.5, MCF7*shMAGI1* untreated EYM= 321.1 +/- 107.1 and treated with Y-27632 EYM= 103.3 +/-79.9. Anova and unpaired two-tailed student's *t*-test; * $p<0.05$ (n=67, 49, 57 and 63 for untreated and treated MCF7*shLuc* and MCF7*shMAGI1*, respectively).

#### SUPPLEMENTARY METHODS

##### Plasmids, mutant constructs and shRNA cloning (Gateway destination vectors)

All the Gateway destination vectors that were generated are listed as follows: *pCMV10 3xFlag RfB-MAGI1, pCMV10 3xFlag RfB-MAGI1 P331A, pCMV10 3xFlag RfB-MAGI1 P390A* and *pCMV10 3xFlag RfB-MAGI1-P331/390A*.

##### Plasmids, mutant constructs and shRNA cloning (Targeted sequences)

*shRNA-MAGI1(1-1):* CACCTATGAAGGAAACTATT
*shRNA-MAGI1(3-1):* GATCTCATAGTGGAAGTTAA
*shRNA-AMOTL2(1416):* GGAACAAGATGGACAGTGA
*shRNA-AMOTL2(3667):* GAGATGTCTTGTTAGCATA
All hairpins were validated (http://cancan.cshl.edu/cgi-bin/Codex/Codex.cgi). The shRNA targeting Luciferase as a control was kindly provided by Dr C. Gongora (IRCM). Retroviral particles were produced in HEK293 cells and used to infect MCF7 and T47D (and HCT116) cell lines that were then selected with 1 µg/ml puromycin.

##### Cell culture

MCF7 cell lines were grown respectively in DMEM:HAM F12 or DMEM medium supplemented with penicillin and streptomycin antibiotics (1%) and fetal calf serum (2% or 10%). T47D and HCT116 were grown in RPMI medium supplemented with antibiotics (1%) and fetal calf serum (2 % or 10%). MCF7, T47D and HCT116 shRNA cell lines were grown in their respective medium in the presence of puromycin (1 µg/mL) for selection of cells expressing shRNAs.

##### Western blotting (antibodies' listing)

Antibodies used for Western blotting were mouse anti-MAG1 (1/250; Santa Cruz #sc100326), rabbit anti-MAGI1 (1/1000; Sigma #HPA031853), rabbit anti-MAGI2 (1/250; Santa Cruz #sc25664), rabbit anti-MAGI3 (1/250; Santa Cruz discontinued), mouse anti-tubulin (1/10000; Sigma-Aldrich #T6074), mouse anti-AMOT (1/250; Santa Cruz #sc166924), rabbit AMOTL2 (1/600; Proteintech #23351-1-AP), mouse mouse anti-GAPDH (1/3000; Proteintech #60004), M2 anti-Flag (1/2000; Sigma-Aldrich #F1804), mouse anti-HA (1/2000; BioLegend #901501), rabbit anti-GFP (1/2000; Torrey Pines Laboratories #TP401), rabbit anti-YAP/TAZ (1/1000; Cell Signaling Technology #8418), rabbit anti YAP (1/1000; Cell Signaling Technology #14074), p-YAP (S127) (1/1000; Cell Signaling Technology #13008) and p-YAP (S397) (1/1000; Cell Signaling Technology #13619), rabbit anti LATS1 (1/1000; Cell Signaling Technology #3477), p-LATS1 (1/1000; Cell Signaling Technology #8654), p-Mob1 (1/1000; Cell Signaling Technology #8699)-mouse anti-actin (1/250; Developmental Studies Hybridoma Bank #JLA20), goat anti-SCRIB (1/250; Santa Cruz #sc11049), rabbit anti-Claudin-1 (1/1000; Cell Signaling Technology #13255), anti-Claudin-3 (1/1000; Genetex #15102), anti-ZO-1 (1/1000; Cell Signaling Technology #8193), anti-β-catenin (1/1000; Cell Signaling Technology #8480), anti-PARD3 (1/1000; Millipore #07-330), anti-E-cadherin (1/1000; Cell Signaling Technology #3195), anti-N-cadherin (1/1000; Cell Signaling Technology #13116), anti-vimentin (1/1000; Cell Signaling Technology #5741), anti-Slug (1/1000; Cell Signaling Technology #9585), anti-Snail (1/1000; Cell Signaling Technology #3879), anti-p-p38 (Cell Signaling Technology #4511), anti-p38 (Cell Signaling Technology #8690), anti-p-JNK/SAPK (Cell Signaling Technology #9251), anti-JNK/SAPK (Cell Signaling Technology #9252), anti-p-MLC Ser19 (Cell Signaling Technology #3675), anti-MLC (Cell Signaling Technology #3672), anti-p-Akt (Cell Signaling Technology #4060), anti-Akt (Cell Signaling Technology #4691), anti-p-ERK1/2 (Cell Signaling Technology #4370) and anti-ERK1/2 (Cell Signaling Technology #4695).

**RNA extraction, reverse transcription and real time RT-qPCR (5’ to 3’ primers listing)**

MAGI1_For: CGTAAAGTGGTTTTTGCGGTGC
MAGI1_Rev: TCTCCACGTCGTAGGGCTGC
MAGI2_For: ATCATTGGTGGAGACGAGCC
MAGI2-Rev: TAGCCACGACACAACACCAG
MAGI3_For: CTGCACTTTTCAGTCTTCTTTTGAC
MAGI3_Rev: CTGAACCAAATTACGTGGCCC
AMOTL2_For: CCAAGTCGGTGCCATCTGTT
AMOTL2_Rev: CCATCTCTGCTCCCGTGTTT
CTGF_For: TTCCAAGACCTGTGGGAT
CTGF_Rev: GTGCAGCCAGAAAGCTC
CYR61_For: ACCAAGAAATCCCCCGAACC
CYR61_Rev: CGGGCAGTTGTAGTTGCATT
BIRC2_For: GTCAGAACACCGGAGGCATT

BIRC2_Rev: TGACATCATCATTGCGACCCA

AREG_For: CGAAGGACCAATGAGAGCCC

AREG_Rev: AGGCATTTCACTCACAGGGG

HPRT_For: CTGACCTGCTGGATTACA

HPRT_Rev: GCGACCTTGACCATCTTT

AXL_For: GTGCCCCTCTCCTTCTTAGC

AXL_Rev: TATGTGACCTGAGCCCCTCT

ATF3_For: CCTCTGCGCTGGAATCAGTC

ATF3_Rev: TTCTTTCTCGTCGCCTCTTTTT

SOX9_For: GAGACTTCTGAACGAGAGCGA

SOX9_Rev: GCCTTGAAGATGGCGTTGG

GATA6_For: CGGGTCAAGATGGGCTCTAC

GATA6_Rev: TGCACAAAAGCAGACACGAG

SOX2_For: ACATGAACGGCTGGAGCAA

SOX2_Rev: GTAGGACATGCTGTAGGTGGG

Axin2_For: GGGGTTGTGTTGGATGGGAT

Axin2_Rev: ATTTCCACGAAAGCACAGCG

CCND1_For: TGGCTGAAGTCACCTCTTGG

CCDN1_Rev: AGCGTATCGTAGGAGTGGGA

p38 alpha_For: TATGGCTCTGTGTGTGCTGCT

p38 alpha_Rev: AAACGTCCAACAGACCAATCAC

**Patients and TMA construction**

The tissue micro-array (TMA) was constructed to encompass the four subtypes of breast cancer. Fifty samples were identified: 14 with hormone receptor positive expression (>10% of tumor cells expressing ER and PR) and HER2-negative (scored 0 by immunohistochemistry), 12 with hormone receptor positive but HER2-positive (scored 3+ by immunohistochemistry), 10 hormone receptor negative HER2 positive and 14 triple negative samples (ER-,PR- and HER2-negative). Tumor samples were collected following French laws under the supervision of an investigator and declared to the French Ministry of Higher Education and Research (Biobanque BB-0033-00059; declaration number DC-2008–695). All patients were informed about the use of their tissue samples for biological research and the study was approved by the local translational research committee.

Tissue blocks with enough material upon gross inspection were initially selected and then the presence of breast carcinoma was evaluated on hematoxylin–eosin-stained sections. The areas to be used for the construction of the TMA were marked on the slide and on the donor block. Samples corresponding to the selected areas were extracted using a manual arraying instrument (Manual Tissue Arrayer 1, Beecher Instruments, Sun Prairie, WI, USA). To take into account the tumor heterogeneity, tumor sampling consisted of two cores (1 mm in diameter) from different tumor areas from a single specimen, and placed at the specified coordinates.

**BrdU incorporation and staining**

One million of cells were incubated with BrdU (50 mg/mL) during 1h30 before being fixed with ethanol 70% at -20°C overnight. Cells were then treated with HCl 2N / Triton-X100 0,5 % and incubated during 30 min at room temperature in the dark. Cells were washed and 1 mL of Borax (pH 8,5) was added before centrifugation. Cells were incubated with anti-BrdU (1/100) during 1h at room temperature in the dark. After washing, cells were incubated with the secondary anti-mouse FITC antibody (1/300) for 1h. Finally, the fluorescent intercalant 7-Aminoactinomycin D (1 µg/mL), along with RNase A, were added to the cells prior to acquirement on a Gallios Flow Cytometer (Beckman Coulter).

**Immunohistochemistry**

PT-Link® system (Dako) was used for pre-treatment, allowing simultaneous de-paraffinization and antigen retrieval. Heat-induced antigen retrieval was executed for 15 minutes in High pH Buffer (Dako) at 95°C. Immunohistochemistry procedure was performed using the Dako Autostainer Link48 platform. Briefly, endogeneous peroxidase was quenched using Flex Peroxidase Block (Dako) for 5 min at room temperature. TMA sections were incubated with rabbit polyclonal antibodies against MAGI1 (1/50; Sigma Aldrich #HPA0311853) and YAP (1/100; Santa Cruz #15407) at room temperature. Following an amplification step with a rabbit linker (Dako) and two rinses in buffer, the slides were incubated with a horseradish peroxidase-labeled polymer coupled to secondary anti-mouse and anti-rabbit antibodies for 20 min, followed by appliance of 3,3'-Diaminobenzidine for 10 min as substrate. Counterstaining was performed using Flex Hematoxylin (Dako) followed by washing the slides under tap water for 5 min. Finally, slides were mounted with a coverslip after dehydration. MCF7 cell line transfected with Flag-MAGI1 expression vector was used as positive control for MAGI1 IHC.

Additional immunohistochemical stainings of primary tumors issued from subcutaneous engraftment of MCF7*shRNA* cells were performed using the Ventana DISCOVERY ULTRA instrument (Ventana Medical Systems, Inc.) and the antibodies used were anti-CD45 (1/500; Bioscience #14-0451-82), anti-Ki67 (1/250; Bioscience #M3064), anti-CD31 (1/75; Abcam #Ab28364) and anti-$\alpha$-SMA (0.02 µg/mL; Ventana #760.2833).

**Immunofluorescence (antibodies' listing)**

Antibodies used were mouse anti-MAGI1 (1/100; Sigma Aldrich #HPA031853), rabbit anti-AMOT (1/100; Proteintech #24550-I-AP), rabbit anti AMOTL2 (1/100; Atlas Antibodies #HPA063027), rabbit anti-E-cadherin (1/100; Cell Signaling Technology #3195), mouse anti-E-cadherin (1/200; BD Transduction Lab #610182), phalloidin (1/200; Sigma-Aldrich #P1951), anti-p-MLC Ser19 (Cell Signaling Technology #3675), rabbit anti-$\beta$-catenin (1/1000; Cell Signaling Technology #8480), rabbit anti-Claudin-3 (1/100; Genetex #GTX15102), anti-PARD3 (1/1000; Millipore #07-330) and anti-ZO-1 (1/100; Cell Signaling Technology #8193) and mouse anti-BrdU (1/100; Santa Cruz #sc32323).

**Thunder Z axis:**

To aquire Z axis images of E-cadherin in our MCF7 cells, we have specifically used an epifluorescence microscope with the Thunder Imaging System (63X objective; Leica Microsystems). The Z axis were generated using the LAS X Life Science microscope software platform.

**Atomic Force Microscopy (AFM) measurements**

Calibrated springs constant for cantilevers were evaluated in the range of 0.15-0.24 N/m. AFM-FS indentation cycles were performed using a 5-10nN force threshold to induce 2-3µm (approx. 20% of cell height) maximal indentation lengths. A squared grid of 4x4 pixels covering a region of 50µm X 50µm was fixed at the center of cell cluster monolayers, defining a force map constituted by 16 indentation curves.

**Mammosphere acquisition and analysis**

At day 5, mammospheres were labeled using 0.2 nM Calcein-AM (PromoCell GmbH) and incubated 30 minutes at room temperature. Data acquisition was done using Incucyte® (Essen Bioscience,

Sartorius) for brigthfield and green fluorescence. Image analysis was processed with IncuCyteS3® Software using the following parameters: Segmentation [Top Hat, Radius (100µm), Threshold (2 GCU)], Edge split [on, edge sensitivity -55], Hole fill [100µm$^2$].

**Flow cytometry**

In summary, 1 x $10^6$ single MCF7*shMAGI1* and MCF7*shLuc* cells were incubated for 30 min with anti-CD44-BV421 (1/200; BioLegend #338809) and CD24-FITC (1/100; BioLegend #101805) antibodies and finally fixed in PBS containing 1 % paraformaldehyde. Aside, the ALDEFLUOR® kit (Stemcell Technologies #01700) was used to isolate the cell population with a high ALDH enzymatic activity according to the manufacturer's conditions. The analyses were performed on a Gallios Flow Cytometer (Beckman Coulter). Precise protocols used for all flow cytometry experiments can be found online (http://www.med.umich.edu/wicha-lab/labmanual8.html).

**TOP/FOP luciferase assay**

In a 96 well plate, MCF7 cells were seeded at a density of 20 000 cells per well. Cells were transfected with 1ng Renilla plus 100ng TOP or FOP. After 48h, cells were lysed using the Dual-Glo luciferase assay system (Promega, #E2920) according to the manufacturer instructions. Renilla and luciferase were then read on MicroBeta TriLux liquid scintillation and luminescence counter.

**Wound healing and Boyden chamber assays**

Wound healing migration assay was performed by scratching confluent MCF7 cell layer with a sterile tip. The area of the wound was measured after 16h to assess cell migration and quantified compared to the control cells. For Boyden chamber invasion assay, Transwell permeable supports (6.5mm Insert with 8µm polycarbonate membrane) were used and coated with 100 µl of matrigel (AMSBIO, #3433-010-01) at a final concentration of 300 µg/ml. MCF7 cells were seeded at a density of 50 000 cells/insert in 1% FBS medium. Under the insert, medium containing 10% FBS was added. As a control, wells without inserts were seeded at the same cell density to assess the proliferation rate. After 48h, cells in the inserts were aspirated and the inside of the insert cleaned with a cotton swab. The medium below the insert was aspirated and MTT was added and the plates incubated for 4h at 37°C. Before reading the OD at 570 nm, DMSO was used to solubilize the formazan crystals formed in the presence of MTT.

**SRB proliferation assay**

3000 cells were plated in 96-well plates and plates were stopped at Day 0, Day 2, Day 5 and Day 8 using 10 % of Trichloroacetic acid (TCA; Sigma Aldrich #T4885) for 10 min, then washed 3 times with PBS and stored at 4°C until the end of the experiment. Once the experiment is finished, 50 µl of Sulforhodamine B (SRB; Sigma Aldrich #230162) 0.04% diluted in acetic acid 1% were added for 30 min before extensive washes with acetic acid 1% and drying step overnight at room temperature. Before reading OD at 540 nm, cells were incubated with Tris 10 mM pH 10.5.

#### Suppl Figure S1

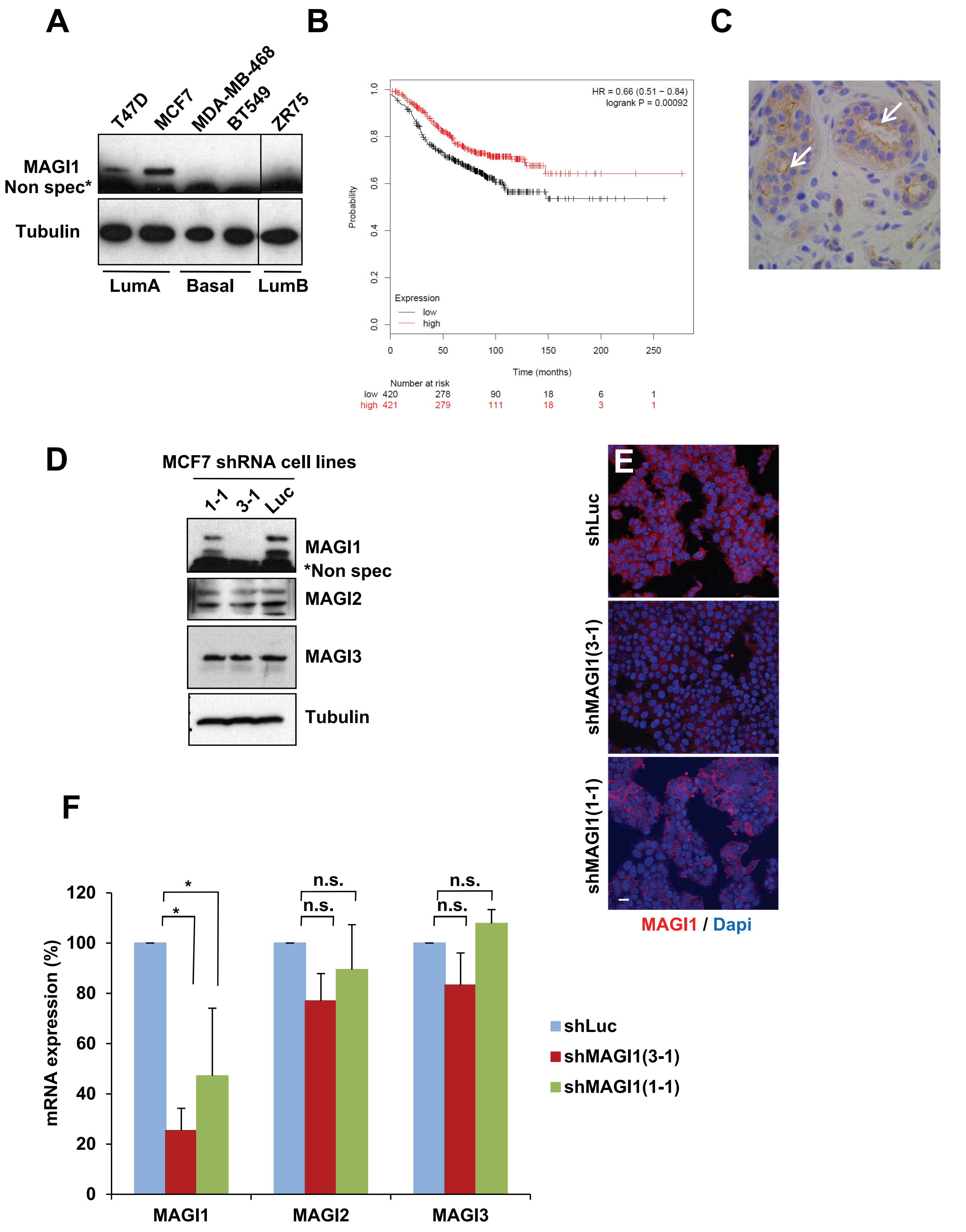

### Suppl Figure S2

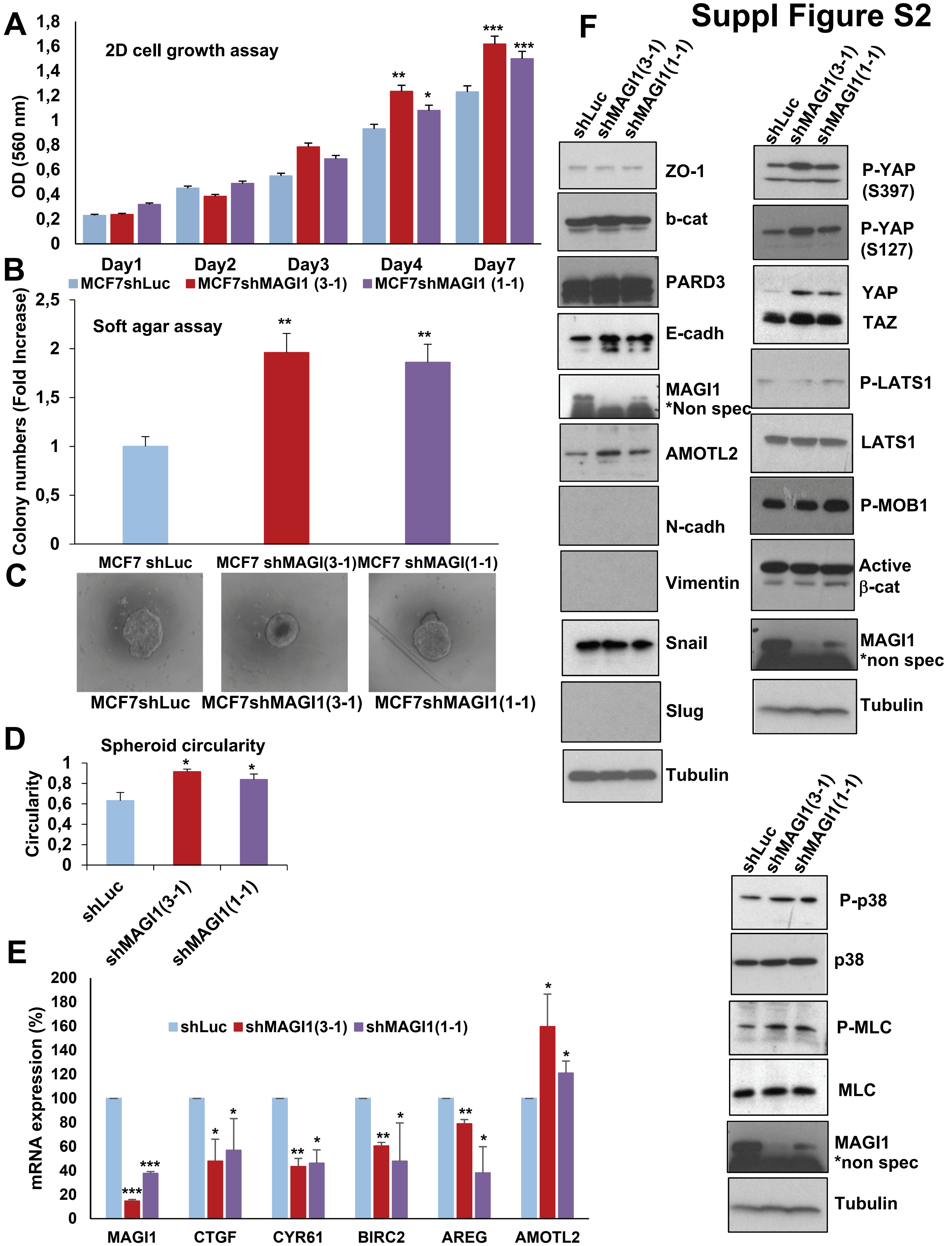

### Suppl Figure S3

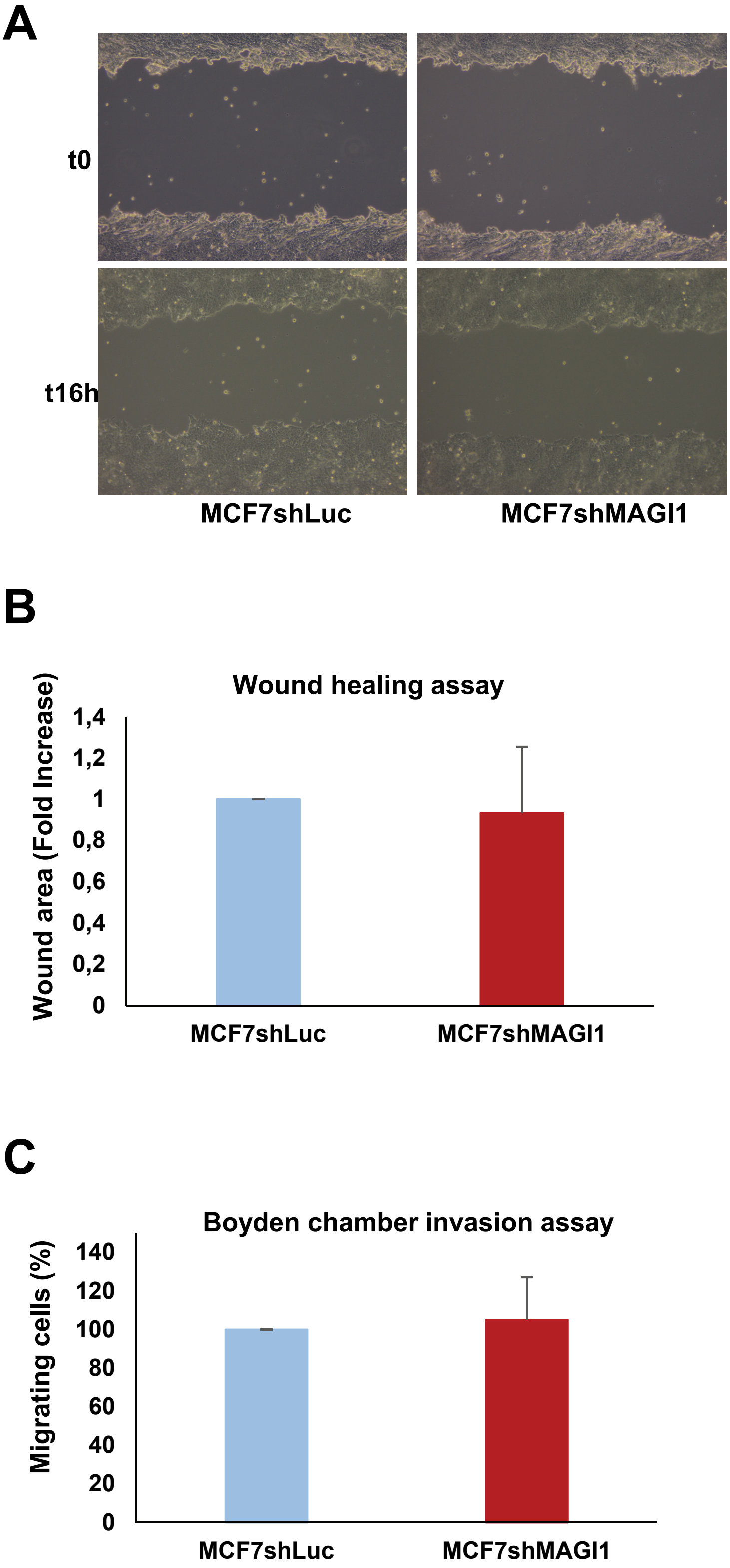

### Suppl Figure S4

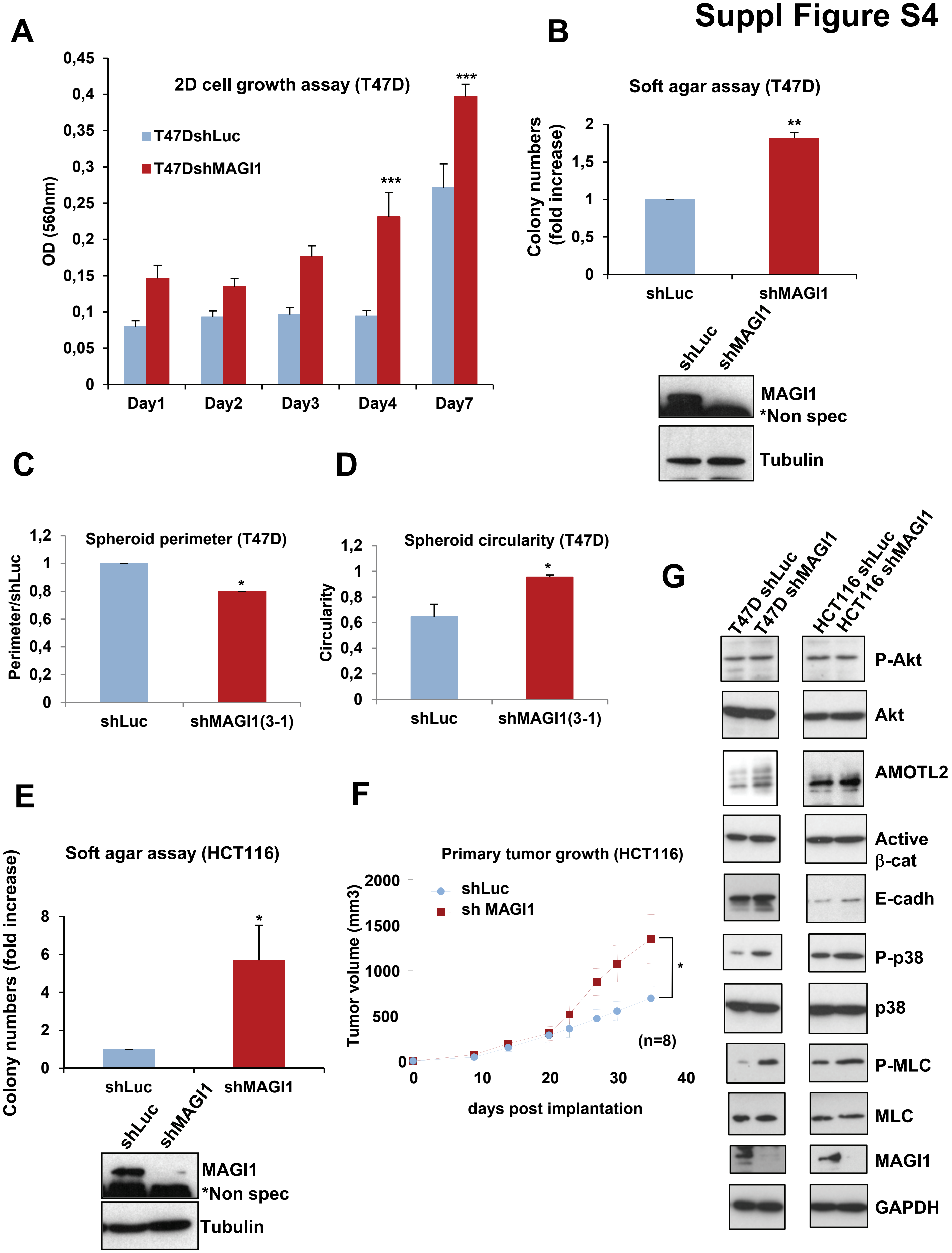

### Suppl Figure S5

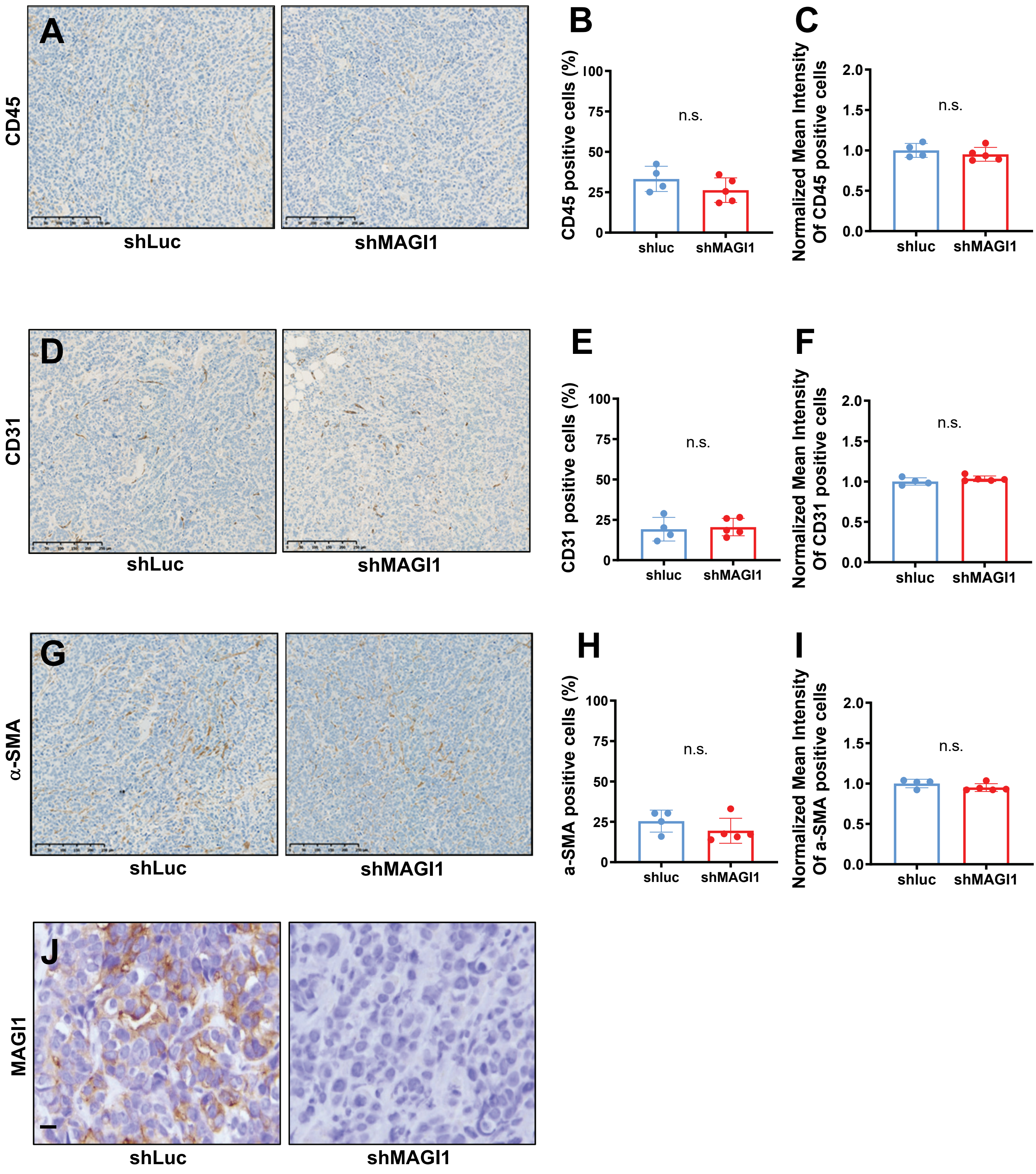

### Suppl Figure S6

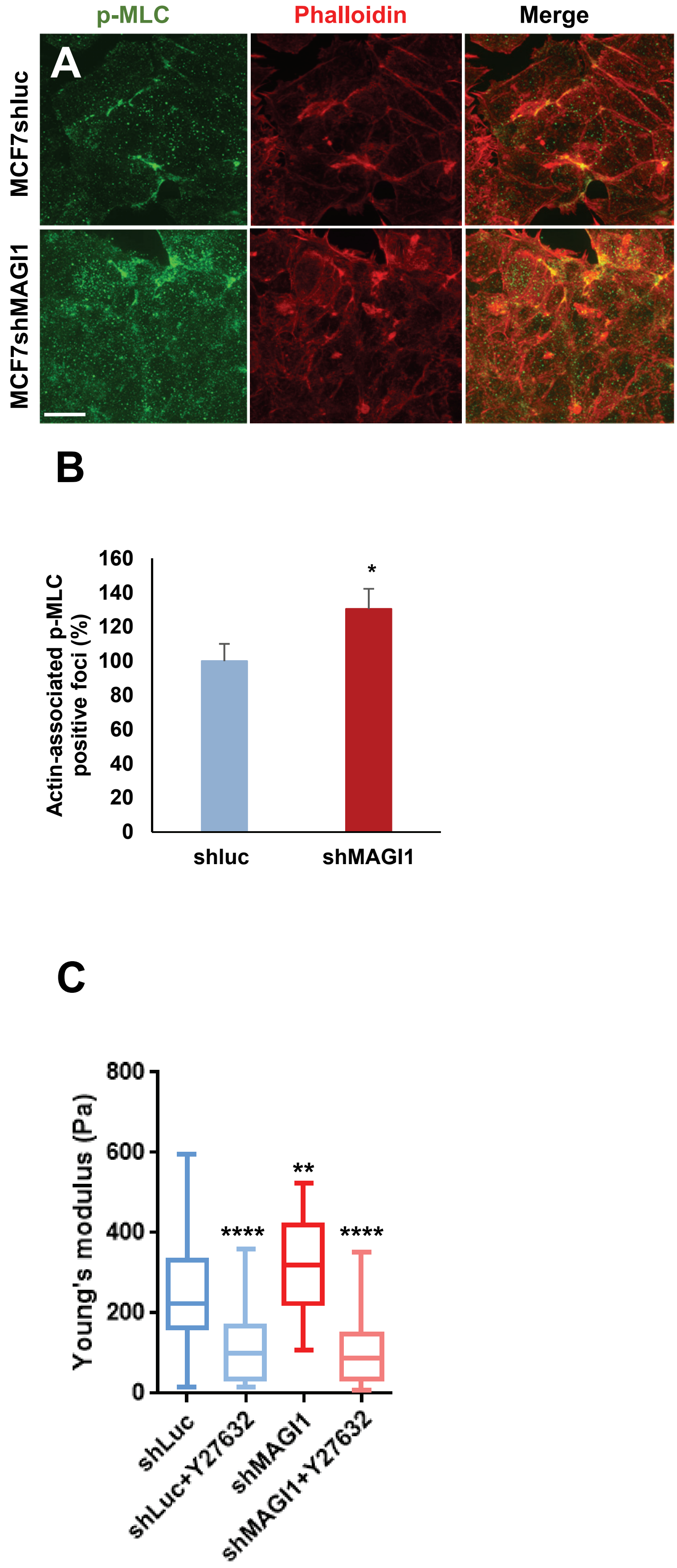

#### Figure 1C

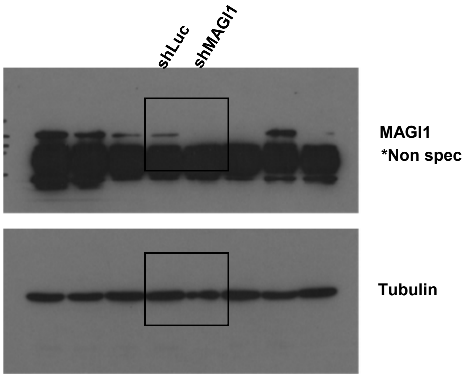


#### Figure 3B

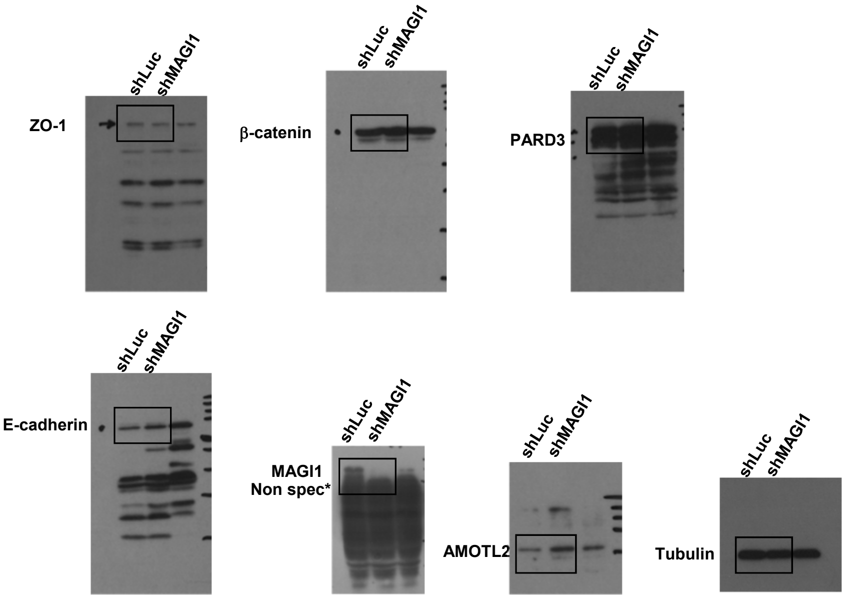


#### Figure 4F

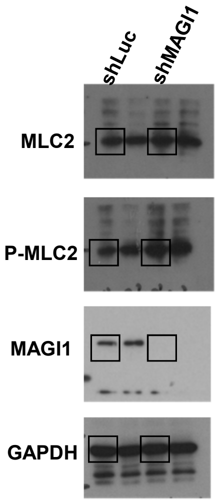


#### Figure 5A

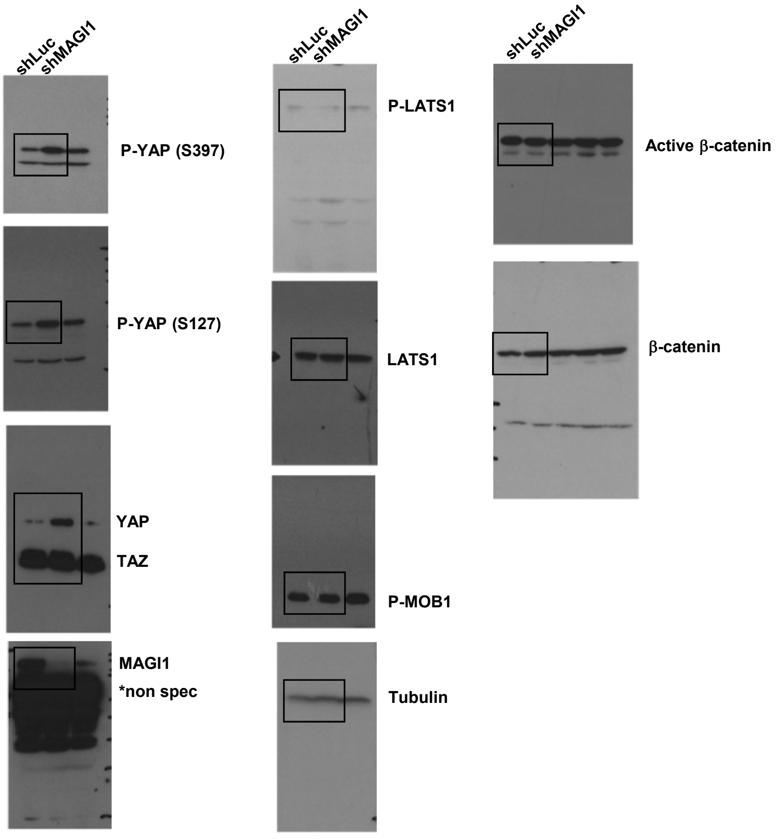

#### Figure 6A

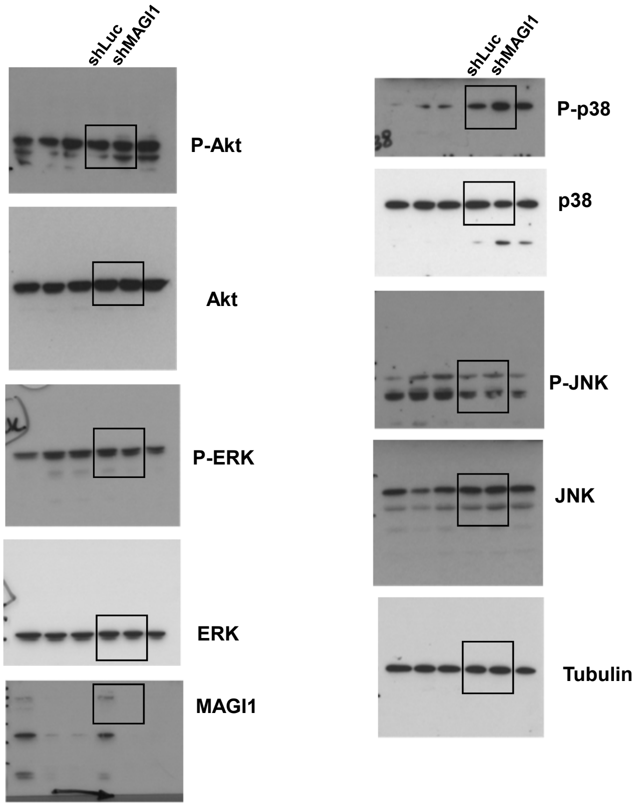


#### Figure 6H

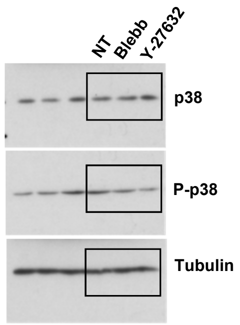


#### Figure 7D

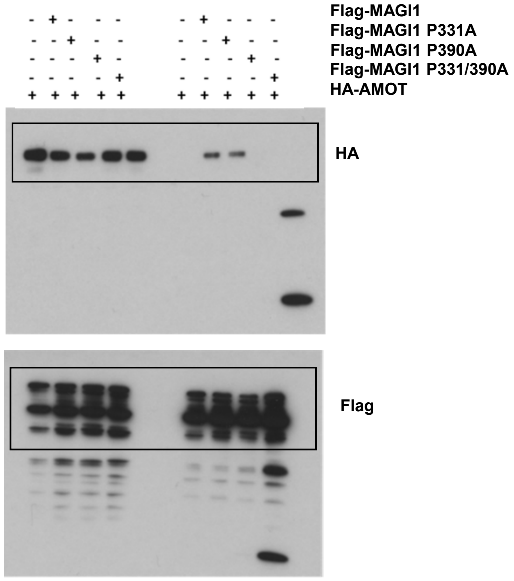


#### Figure 7A

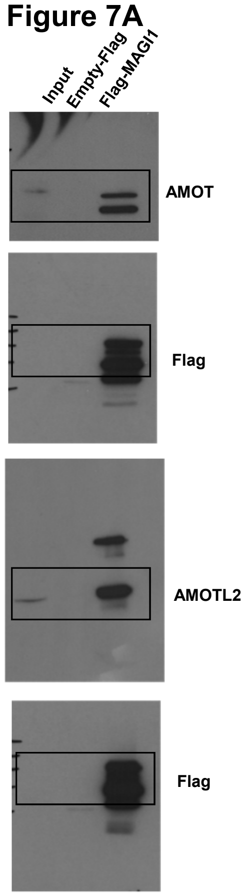


#### Figure 7B

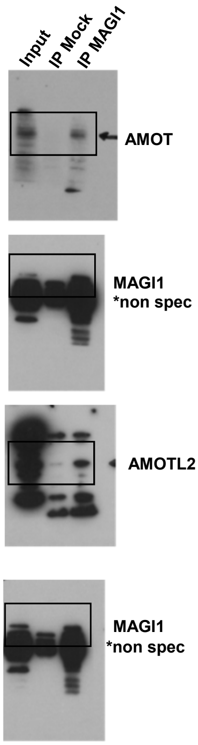


#### Figure 7E

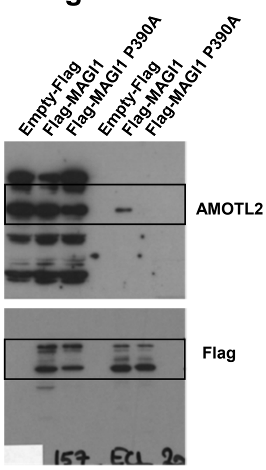


#### Figure 7G

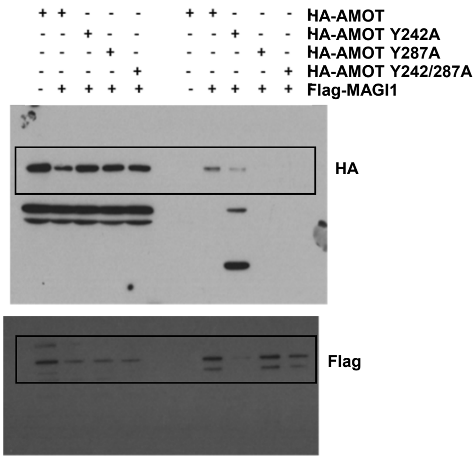

#### Figure 7I

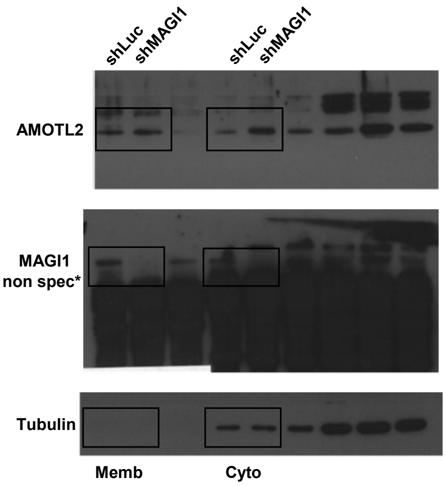


#### Figure 8C

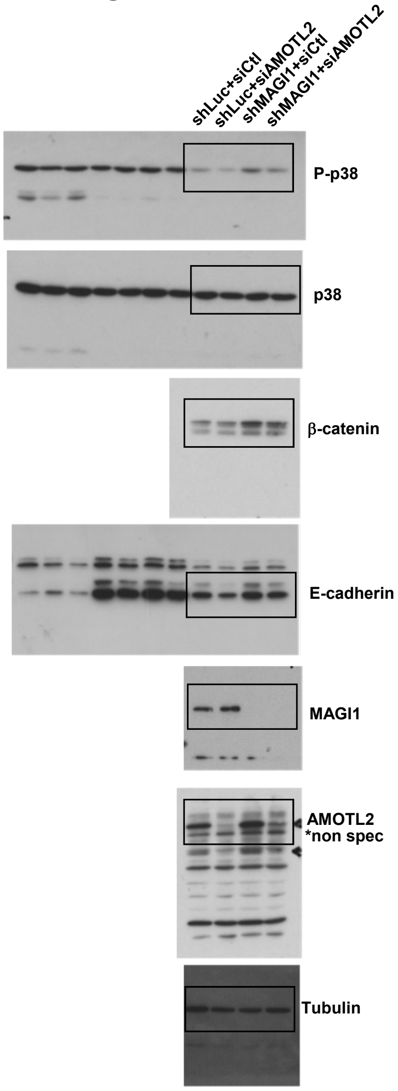


#### Suppl Figure S1A

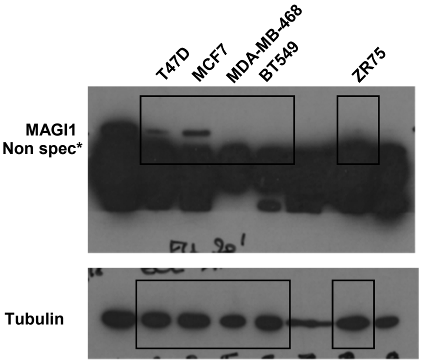


#### Suppl Figure S1D

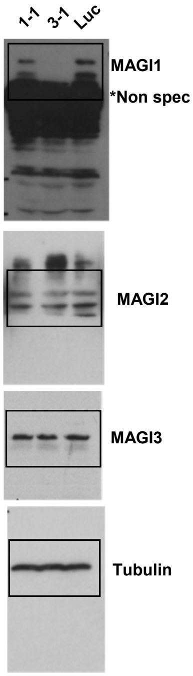

#### Suppl Figure S2F

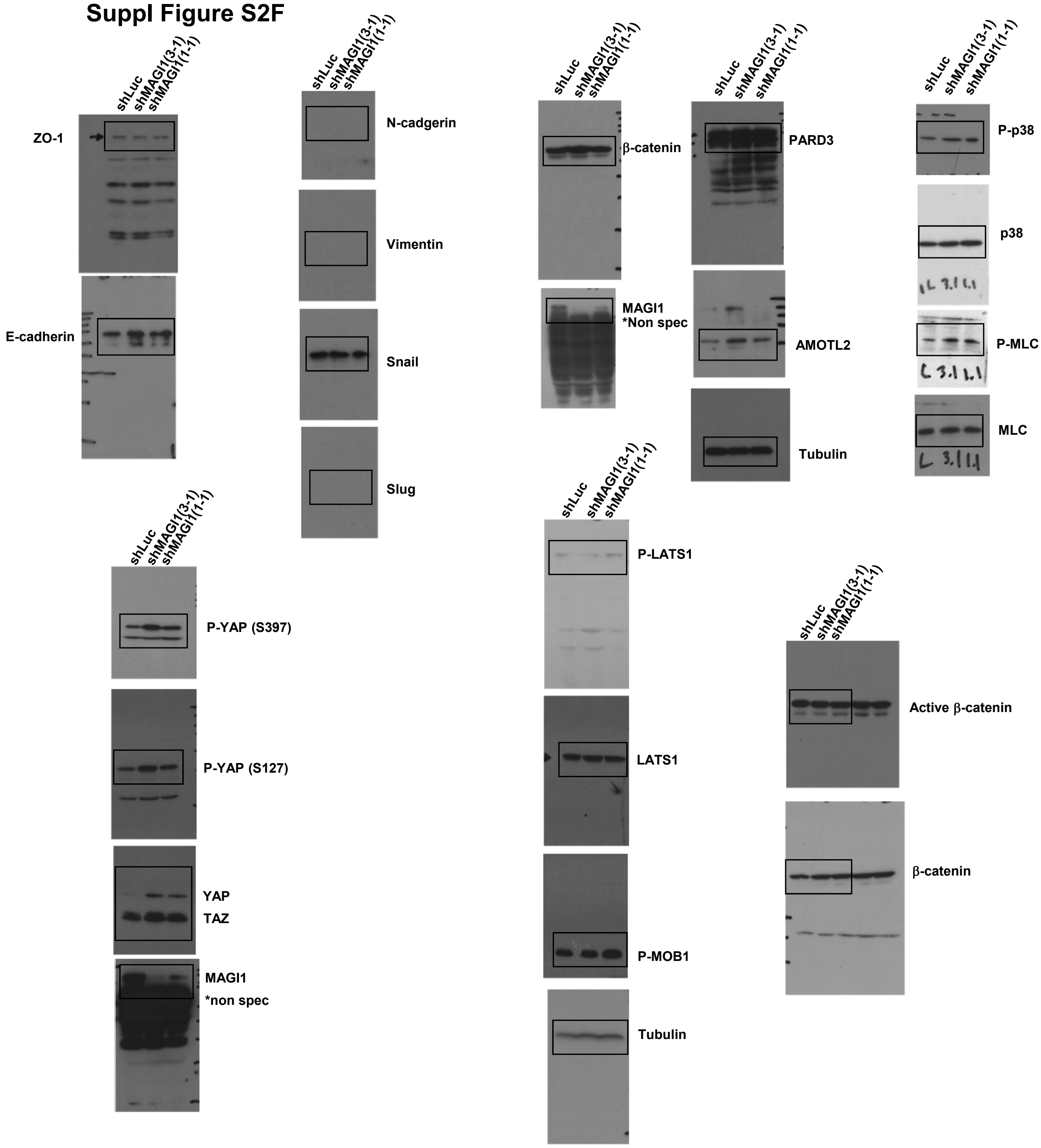


#### Suppl Figure S4B

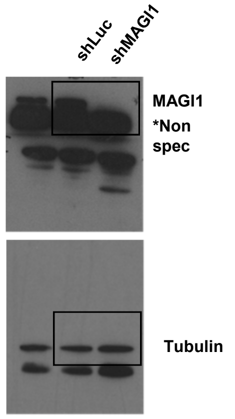


#### Suppl Figure S4E

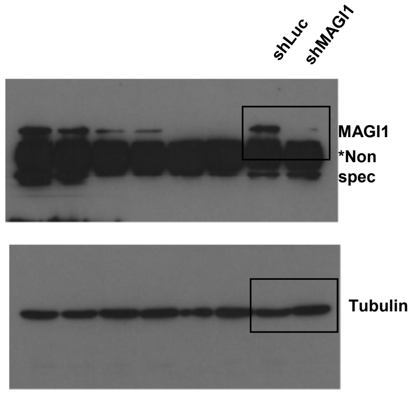

#### Suppl Figure S4G

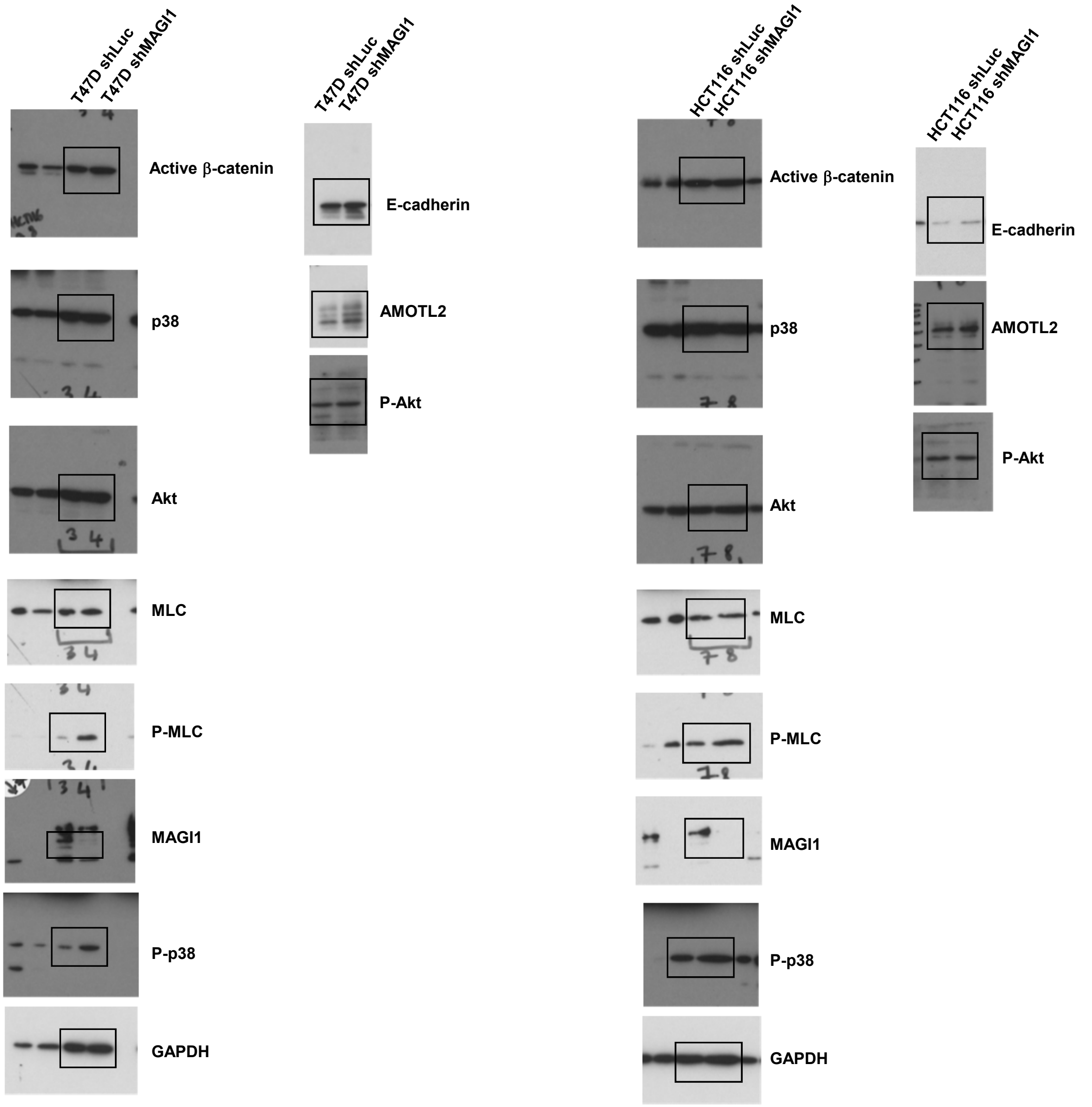
